# β-adrenergic receptor activation modulates the induction of complex spike burst-dependent LTP by regulating multiple forms of heterosynaptic plasticity

**DOI:** 10.1101/2025.04.28.651102

**Authors:** Thomas J. O’Dell

**Author notes:** Correspondence: T. J. O’Dell, Department of Physiology, David Geffen School of Medicine at UCLA, 53-231 Center for the Health Sciences, Box 951751, Los Angeles, CA 90095-1751.

## Abstract

The modulatory neurotransmitter norepinephrine and activation of β-adrenergic receptors (β-ARs) has a crucial role in hippocampus-dependent forms of learning. Although β-AR activation also facilitates the induction of Hebbian LTP at excitatory synapses in the hippocampus, recent findings indicate that a non-Hebbian form of synaptic plasticity, known as behavioral timescale synaptic plasticity (BTSP), underlies hippocampus-dependent spatial learning. Because little is known about the role of noradrenergic signaling in BTSP, I examined the effects of the β-AR activation on a BTSP-like, complex spike (CS) burst-dependent form of LTP induced by theta-pulse stimulation (TPS) protocols in the CA1 region of mouse hippocampal slices. I find that β-AR activation not only enhances the homosynaptic potentiation of synaptic transmission induced by TPS but also regulates how synapses interact during the induction of CS burst-dependent LTP. The ability of synapses to interact in a cooperative fashion to undergo CS burst-dependent LTP, which is mediated by a heterosynaptic facilitation of EPSP-evoked CS bursting induced by brief trains of TPS, was enhanced by β-AR activation. Moreover, unlike conventional Hebbian LTP, where cooperative LTP induction requires near synchronous coactivation of synapses, β-AR activation enabled the induction of CS burst-dependent LTP by cooperative interactions between synapses activated up to ten seconds apart. β-AR activation also enhanced a potent form of synaptic competition that emerges during longer trains of TPS. This effect of β-AR activation is mediated by a CS burst-dependent form of heterosynaptic depression and produces an unusual, bidirectional modulation of LTP induction where the β-AR-mediated facilitation of homosynaptic LTP induction produces a heterosynaptic suppression of LTP induction at other synapses. Together, these findings indicate that β-AR activation regulates fundamental properties of postsynaptic burst-dependent plasticity rules by modulating multiple forms of heterosynaptic plasticity.

## 1 INTRODUCTION

The modulatory neurotransmitter norepinephrine (NE), acting through β-adrenergic receptors (β-ARs), enhances learning and memory (Sara, 2009; Hagena et al., 2016) as well as the induction of Hebbian LTP, a form of synaptic plasticity thought to underlie memory formation (O’Dell et al., 2015). Importantly, associative forms of learning are highly supervised and dependent on factors such as continency (i.e., the predictive value of a conditional stimulus), attention, and novelty or prediction errors (Gallistel and Matzel, 2013; Fanselow and Wassum, 2016). In contrast, Hebbian LTP is a correlation-based form of plasticity where increases in synaptic strength can be induced in an unsupervised manner by temporally contiguous pre- and postsynaptic activity. Thus, although considerable evidence supports the notion that Hebbian LTP is involved in memory formation (Dringenberg, 2019), the unsupervised nature of changes in synaptic strength produced by Hebbian LTP is strikingly incompatible with fundamental properties of associative learning (Gallistel and Matzel, 2013). Notably, noradrenergic neurons in the locus coeruleus are activated by novel stimuli (Sara et al., 1994; Takeuchi et al., 2016), changes in stimulus-reward contingencies (Sara and Segal, 1991; Bouret and Sara, 2004), and prediction errors (Jorden, 2024), indicating that these cells encode information that has a crucial role in associative memory formation. The noradrenergic modulation of Hebbian LTP may thus provide a mechanism that allows synapses to use a simple, correlation-based form of plasticity to store information in a more computationally sophisticated and behaviorally relevant manner (Frémaux and Gerstner, 2016).

In-vitro studies demonstrating a modulatory role for NE in synaptic plasticity have primarily used stimulation protocols that induce standard Hebbian LTP, such as bouts of high-frequency synaptic stimulation or repeated pairing of pre- and postsynaptic action potentials (O’Dell et al., 2015; Brzosko et al., 2019). Growing evidence indicates, however, that a non-Hebbian form of LTP, known as behavioral timescale synaptic plasticity (BTSP), underlies hippocampus-dependent forms of spatial learning (Bittner et al., 2017; Priestley et al., 2022; Grienberger and Magee, 2022; Zhao et al., 2022). In BTSP, synaptic inputs that trigger dendritic plateau potentials act as an instructive signal that enables LTP induction at other synapses activated seconds before or after plateau potentials (Bittner et al., 2017; Xiao et al., 2023). Notably, dendritic plateau potentials elicit somatic complex-spike (CS) bursts in CA1 pyramidal cells (Takahashi and Magee, 2009; Grienberger et al., 2014) and a CS burst-dependent form of LTP in CA1 pyramidal cells has non-Hebbian properties like those seen in BTSP (O’Dell, 2022). Given the information encoded by neurons in the locus coeruleus, NE likely has an important role in both BTSP and CS burst-dependent LTP. Consistent with this notion, activation of locus coeruleus axons in the hippocampus has a crucial role in generating place-specific firing in CA1 pyramidal cells when animals encounter unexpected rewards in a familiar environment, presumably through modulation of BTSP (Kaufman et al., 2022). Moreover, β-AR activation facilitates the induction of CS burst-dependent LTP (Thomas et al., 1996; Winder et al., 1999). Beyond this, however, little is known about the effects of β-AR activation on the properties of BTSP and CS burst-dependent LTP.

Although EPSP-evoked CS bursts elicited during theta-frequency trains of synaptic stimulation induce a Hebbian form of homosynaptic LTP (Thomas et al., 1998), the induction of CS burst-dependent LTP is also strongly dependent on non-Hebbian forms of heterosynaptic plasticity (O’Dell, 2022). For example, brief bouts of EPSP-evoked CS bursting that induce relatively modest amounts of homosynaptic LTP also induce a short-term, heterosynaptic facilitation of CS bursting that enables asynchronously activated synapses to interact in a cooperative fashion and undergo LTP (O’Dell, 2022). In contrast, longer bouts of EPSP-evoked CS bursting that induce more robust homosynaptic LTP trigger a form of synaptic competition that inhibits LTP induction at other synapses (O’Dell, 2022). Given the crucial role of cooperativity and competition in CS burst-dependent LTP, the heterosynaptic forms of plasticity underlying these processes may provide important targets where NE can act to regulate CS burst-dependent changes in synaptic strength. Thus, in this study I investigated the effects of β-AR activation on synaptic interactions during the induction of CS burst-dependent LTP.

## 2 METHODS

### 2.1 Animals and Slice Preparation

Hippocampal slices from the dorsal hippocampus were obtained from 8-14 weeks old, male and female C57Bl/6 mice (#027 Charles River Laboratories). Mice were deeply anesthetized with isoflurane and, following cervical dislocation, the brain was removed and place in cold (approximately 4°C), oxygenated (95% O_2_/5% CO_2_) ACSF containing 124 mM NaCl, 4 mM KCl, 25 mM NaHCO_3_, 1 mM NaH_2_PO_4_, 2 mM CaCl_2_, 1.2 mM MgSO_4_, and 10 mM glucose. Hippocampi were then dissected from the brain and 400-µm-thick slices were prepared using a manual tissue slicer. The CA3 region was removed, and slices were then transferred into interface-type chambers continuously perfused with ACSF (2-3 mL/min) and allowed to recover (at 30°C) for at least 2 hours before recordings. All techniques were approved by the Institutional Animal Care and Use Committee at the University of California, Los Angeles and done in compliance with U.S. Public Health Service guidelines.

### 2.2 Electrophysiological Recordings

Extracellular recordings were done using slices maintained at 30°C in an interface-type recording chamber perfused with ACSF. Two bipolar, nickel-chromium wire stimulating electrodes were placed in stratum radiatum to activate independent groups of Schaffer collateral/commissural fiber synapses onto CA1 pyramidal cells (hereafter referred to as S1 and S2 synapses). A glass microelectrode filled with ACSF (resistance ≍ 10 MΩ) was placed in stratum radiatum to record field excitatory postsynaptic potentials (fEPSPs). Signals were acquired with a Multi-Clamp 700B amplifier (Molecular Devices), low pass filtered at 2 kHz, and digitized at 10 kHz. fEPSPs were evoked by capacity-coupled, constant-voltage pulses (0.02 msec duration) delivered by a Grass S88 stimulator. After determining the maximal amplitude of fEPSPs evoked by each stimulating electrode, the stimulation intensity was adjusted to evoke fEPSPs with an amplitude approximately 50% of the maximal amplitude. Standard techniques were used to ensure activation of independent groups of synapses (O’Dell, 2022) and alternating pulses of S1 and S2 presynaptic fiber stimulation were then delivered at 0.04 Hz. All chemicals were obtained from Millipore-Sigma except for tetrodotoxin (TTX) (Alomone Labs) and gabazine (Abcam). Stock solutions of the β-AR agonist isoproterenol (ISO) were prepared fresh daily (1 mM in dH_2_O). Stock solutions of gabazine (1 mM) and TTX citrate (1 mM) were prepared in dH_2_O and stored at -20 °C. Stock solutions of 8-cyclopentyl-1,3-dipropylxanthine (DPCPX, 2 mM) were prepared in DMSO and stored at -20 °C.

### 2.3 Experimental Design and Statistical Analysis

CS burst-dependent LTP was induced using theta-pulse stimulation (TPS) protocols (single pulses of presynaptic fiber stimulation delivered at 5 Hz). Average slopes of fEPSPs (normalized to baseline) recorded 40-45 minutes post-TPS were used for statistical comparisons. EPSP-evoked CS bursting during TPS was quantified by visually inspecting fEPSPs evoked during TPS and counting the number of negative-going population spikes (PSs) elicited by each EPSP during the train. CS bursts were defined as fEPSPs containing 2 or more PSs. Probability of EPSP-evoked CS bursting (*P*_CSB_), determined from the number of EPSPs eliciting CS bursts relative to total number of EPSPs evoked during TPS, was used for statistical comparisons. Heterosynaptic depression was measured using the slope of the first S2 fEPSP (normalized to baseline) elicited after a train of TPS delivered to S1 synapses. No obvious sex differences were found and the results from male and female mice were combined. Student’s t-tests or, where appropriate, Mann-Whitney Rank Sum tests were used to evaluate statistical significance between two groups. Correlations were determined using Pearson Product Moment Correlation tests. Multiple comparisons were done using one-way or two-way ANOVAs with Student-Newman-Keuls (SNK) *post hoc* comparisons or Kruskal-Wallis one-way ANOVA on Ranks followed by Dunn’s tests for multiple, pairwise comparisons. Data were collected and analyzed using pClamp10 software (Molecular Devices). Statistical tests were performed using SigmaPlot 12.5 (Grafiti). Results are reported as mean ± SEM and full results of statistical tests are provided in the figure legends.

## 3 RESULTS

### 3.1 Cooperative and Competitive Synaptic Interactions Regulate the Induction of CS Burst-Dependent LTP

Results shown in figures 1 and 2 highlight four key properties of CS burst-dependent LTP. First, during TPS the mode of action potential firing in CA1 pyramidal cells gradually transitions from single spike firing to burst firing, leading to an activity-dependent emergence of EPSP-evoked bursting (Fig. 1A). A previous study using somatic whole-cell current-clamp recordings demonstrated that these bursts, which appear as multiple, negative-going PSs in dendritic field potential recordings (Fig. 1A), exhibit many of the defining features of CS bursts recorded in-vivo (O’Dell, 2022). Second, because CS bursts provide the postsynaptic depolarization needed for NMDA receptor (NMDAR) activation during TPS (Thomas et al., 1998; O’Dell, 2022), the induction of LTP by different patterns of TPS reflects the activity-dependence of EPSP-evoked CS bursting. For example, 30-seconds of TPS delivered to S2 synapses elicited robust EPSP-evoked CS bursting (*P_CSB_* = 0.71, n = 10) and induced a synapse-specific potentiation of S2 synapses (fEPSPs potentiated to 156 ± 4% of baseline) (Fig. 1B). In contrast, a brief, 5-seconds-long train of TPS that ended before the onset of CS bursting (*P_CSB_* = 0.07, n = 10) had no lasting effect on synaptic strength (45 minutes post-TPS fEPSPs were 102 ± 3% of baseline) (Fig. 2A). Third, unlike conventional Hebbian induction protocols, where near synchronous coactivation of synaptic inputs is required for synapses to interact in a cooperative fashion and undergo LTP (Gustafsson and Wigström, 1986; Lin et al., 2003), CS burst-dependent LTP can be induced in a highly cooperative manner by sequential, asynchronous activation of independent groups of synapses (O’Dell, 2022). An example of this is shown in figure 2B, where 10 seconds of TPS was delivered to S1 synapses to bring CA1 pyramidal cells to threshold for EPSP-evoked CS bursting before 5 seconds of TPS was delivered to S2 synapses. Although 5 seconds of TPS delivered to S2 synapses by itself elicits few CS bursts (Fig. 2A), EPSP-evoked CS bursting at S2 synapses was strongly enhanced when S2 TPS was delivered after 10 seconds of TPS delivered to S1 synapses (*P_CSB_* during S2 TPS = 1.0, n = 8) and S2 fEPSPs potentiated to 164 ± 4% of baseline (Fig. 2B). Due to the activity dependence of CS bursting during TPS (Fig. 1A), relatively few of the EPSPs evoked during S1 TPS elicited CS bursts (*P_CSB_* = 0.3) and S1 synapses potentiated to just 131 ± 4% of baseline (Fig. 2B). Thus, although 10 seconds of TPS induces relatively modest homosynaptic LTP, it produces a powerful heterosynaptic facilitation of EPSP-evoked bursting and LTP induction at S2 synapses. Indeed, the potentiation of S2 synapses induced by the S1→ S2 TPS protocol used in these experiments was 2-fold larger than that seen at S1 synapses (*t*_(14)_ = 5.353, *p* = 1.02 x 10^-4^), even though the S2 TPS train was only half as long as the TPS train delivered to S1 synapses. Finally, competitive synaptic interactions that emerge during longer trains of TPS also potently regulate the induction of CS burst-dependent LTP (O’Dell, 2022). For example, although increasing the duration of S1 TPS to 20 seconds significantly enhanced homosynaptic LTP at S1 synapses (S1 fEPSPs potentiated to 164 ± 5% of baseline, n = 10), the heterosynaptic facilitation of TPS-induced LTP at S2 synapses was abolished (45 minutes post-TPS S2 fEPSPs were 101 ± 2% of baseline), even though EPSPs elicited CS bursts during S2 TPS (*P_CSB_* = 0.85) (Fig. 2C). Thus, patterns of TPS that induce robust homosynaptic LTP also generate a potent, winner-take-all form of synaptic competition that blocks LTP induction at other synapses (Fig. 2D).

**Figure 1.**
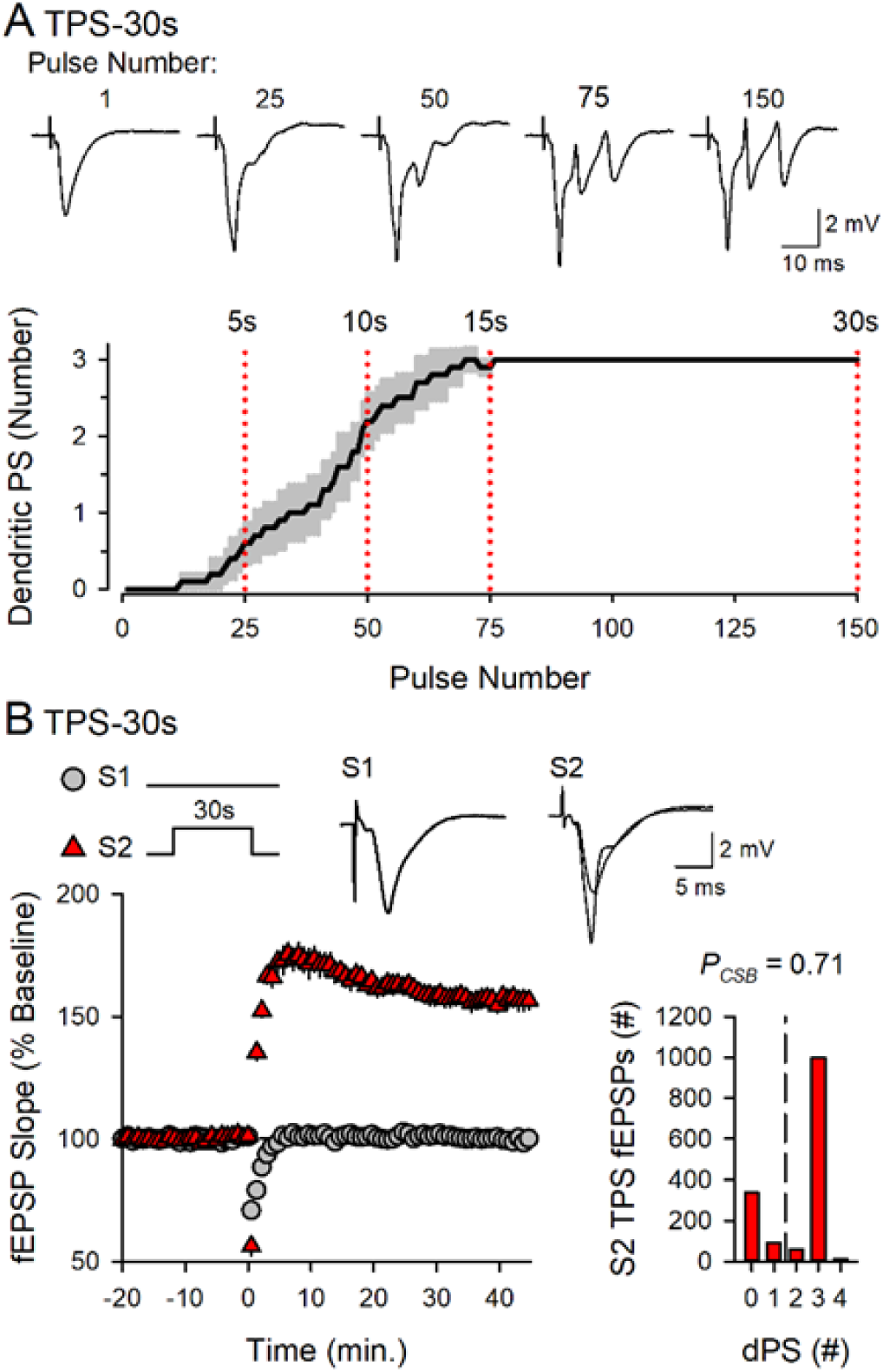
Theta-pulse stimulation (TPS) triggers EPSP-evoked CS bursting and induces LTP. ***A*:** Number of dendritic population spikes (dPS) evoked by EPSPs during a 30-seconds-long train of TPS. Shading represents ± SEM (n = 10). Traces show fEPSPs evoked by the indicated pulse numbers during TPS. ***B*:** LTP induced by 30 seconds of TPS delivered to S2 synapses at time = 0. Results are from the same experiments shown in A. Traces show superimposed fEPSPs elicited by S1 and S2 stimulation during baseline and 45 minutes post TPS. Histogram shows the number of EPSPs evoking 0-4 dPSs during TPS from all experiments.

**Figure 2.**
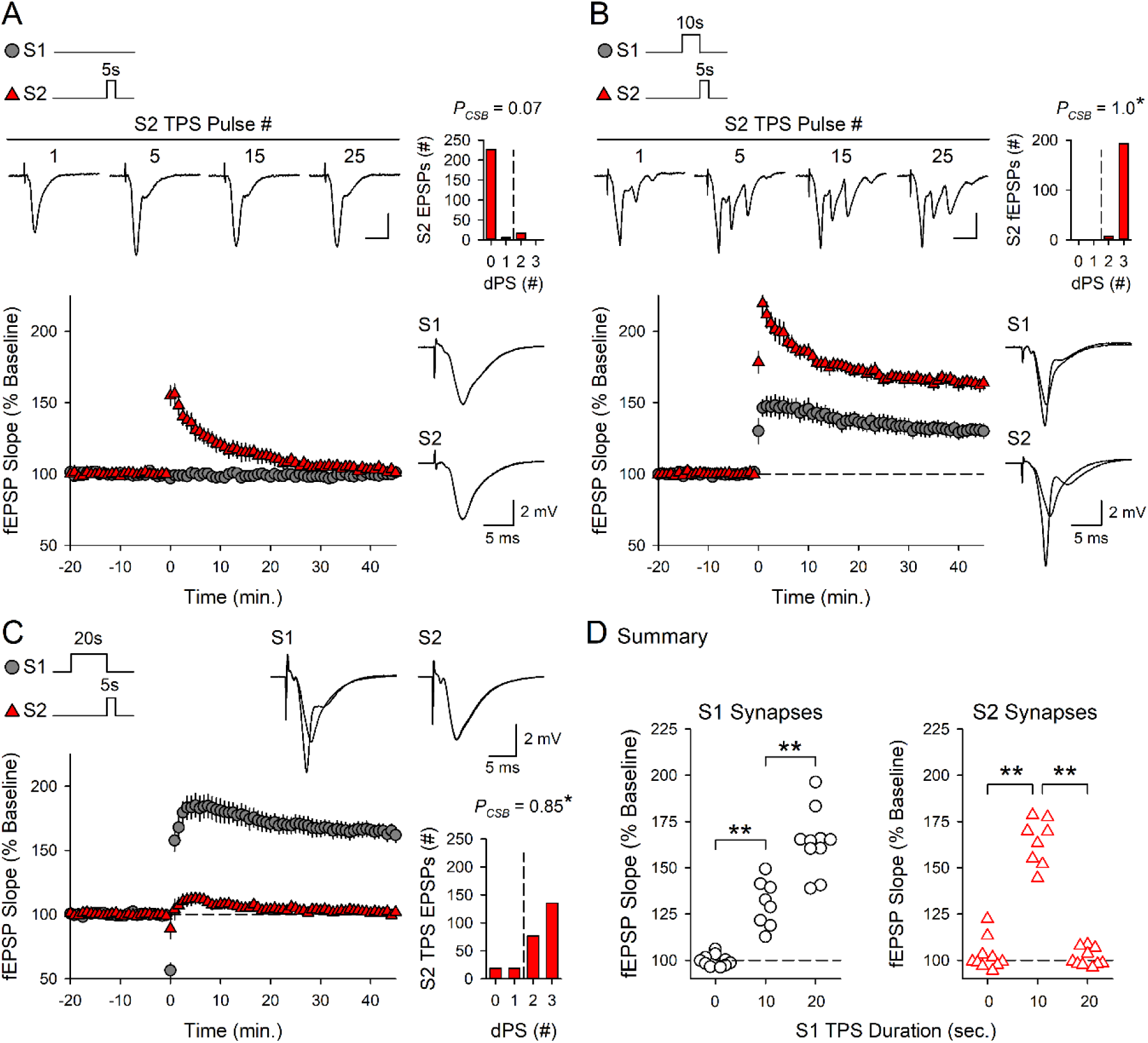
Cooperative and competitive synaptic interactions regulate CS burst-dependent LTP induction. ***A*:** 5 seconds of TPS delivered to S2 synapses alone (at time = 0) failed to elicit CS bursting and had no lasting effect on synaptic strength (n = 10). ***B*:** 10 seconds of TPS delivered to S1 synapses before 5 s of S2 TPS facilitates LTP induction at S2 synapses (n = 8). Traces in A and B show fEPSPs evoked by the indicated pulses during S2 TPS (top, calibration bars are 2 mV and 10 ms) and superimposed fEPSPs elicited by S1 and S2 stimulation during baseline and 45 minutes post-TPS (right). ***C*:** 20 seconds of TPS delivered to S1 synapses before 5 seconds of S2 TPS inhibits LTP induction at S2 synapses (n = 10). Traces show superimposed fEPSPs evoked by S1 and S2 stimulation during baseline and 45 minutes post-S2 TPS. Histograms in *A*-*C* show the number of EPSPs evoking 0-3 dendritic PSs during S2 TPS. \**p* < 0.05 compared to *P_CSB_* during 5 seconds of TPS delivered to S2 synapses alone, one-way ANOVA on Ranks with Dunns *post hoc* tests, *H*_(2)_ = 21.363, *p* < 0.001. ***D*:** S1 (left) and S2 (right) fEPSP slopes 45 minutes after S2 TPS from all experiments in *A*-*C*. Changes in synaptic strength at S1 and S2 synapses were analyzed separately using one-way ANOVAs with SNK *post hoc* comparisons (\*\**p* < 0.001, S1 synapses: *F*_(2,25)_ = 68.107, *p* < 0.001; S2 synapses: *F*_(2,25)_ = 140.264, *p* < 0.001).

### 3.2 β-AR Activation Enhances Competitive Synaptic Interactions During the Induction of CS Burst-Dependent LTP

Numerous studies have found that β-AR activation enhances TPS-induced LTP (Thomas et al., 1996; Winder et al., 1999; Gelinas and Nguyen, 2005; Gelinas et al., 2008; Qian et al., 2012; Jami et al., 2023). Consistent with these findings, although 5 seconds of TPS delivered to S2 synapses had no lasting effect on synaptic transmission in control experiments (Fig. 3A), S2 fEPSPs potentiated to 153 ± 8% of baseline (n = 9) when TPS was delivered at the end of a 10-minute bath application of 1.0 µM ISO (Fig. 3B,C). To explore the effects of β-AR activation on synaptic interactions during the induction of CS burst-dependent LTP, I next examined the effects of ISO on the heterosynaptic facilitation of LTP induction at S2 synapses produced by a 15-seconds-long train of TPS delivered to S1 synapses. In control experiments, 15 seconds of TPS delivered to S1 synapses before a 5-seconds-long train of S2 TPS facilitated EPSP-evoked CS bursting during S2 TPS (*P_CSB_* = 0.99, n = 9) and enabled LTP induction at S2 synapses (S2 fEPSPs potentiated to 149 ± 6% of baseline) (Fig. 3D). Surprisingly, although LTP induction at S1 synapses was enhanced when this pattern of S1→S2 TPS was delivered in the presence of ISO, LTP induction at S2 synapses was strongly suppressed (45 minutes post-TPS in ISO S2 fEPSPs were 113 ± 3% of baseline, n = 10) (Fig. 3E,F). This paradoxical ability of ISO to simultaneously enhance LTP induction at S1 synapses and inhibit the induction of LTP at S2 synapses indicates that β-AR activation modulates the heterosynaptic forms of plasticity underlying synaptic interactions during CS burst-dependent LTP induction.

**Figure 3.**
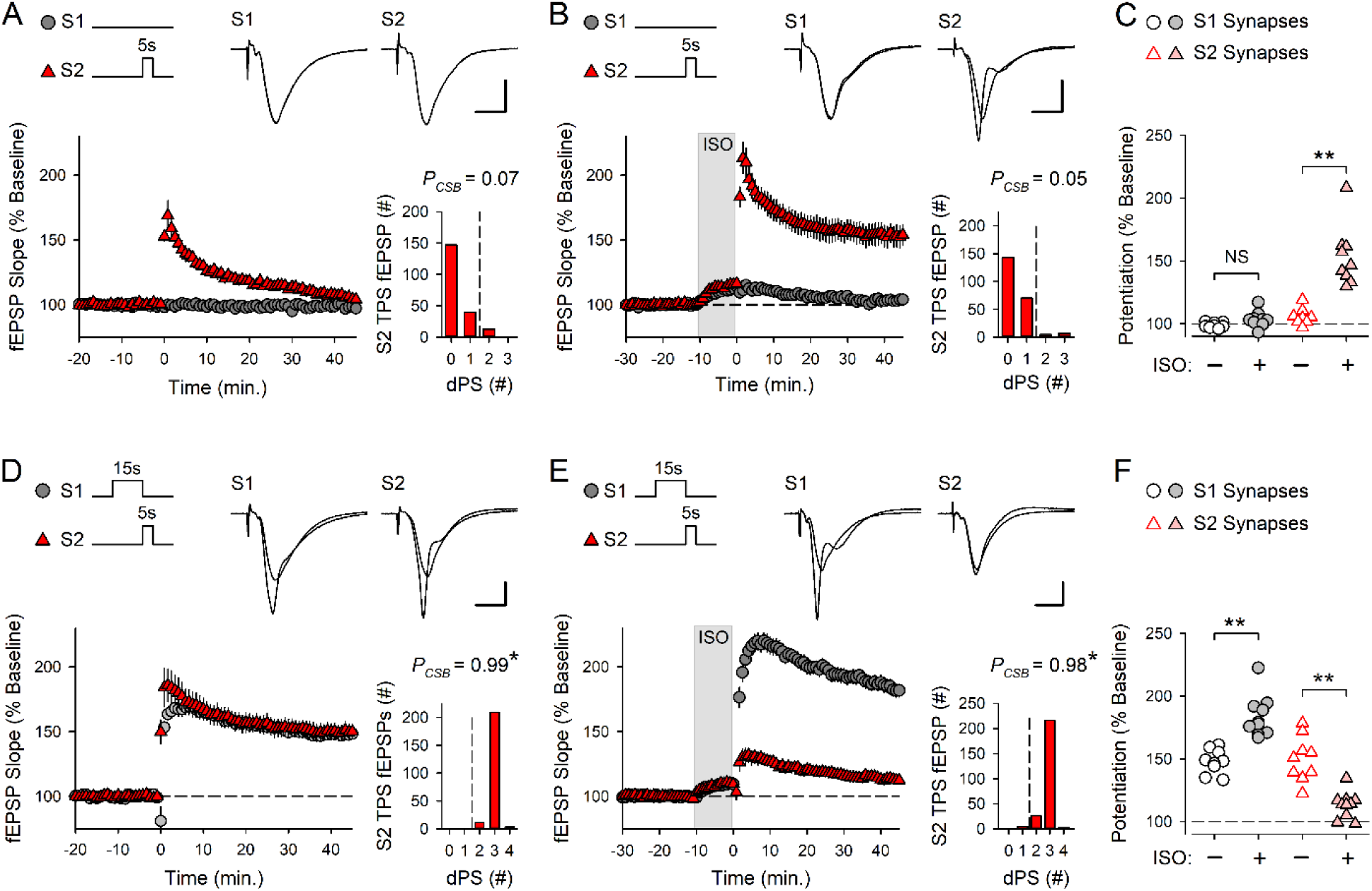
β-AR activation enhances LTP and competitive synaptic interactions during the induction of CS burst-dependent LTP. ***A*:** Control experiments where 5 seconds of TPS was delivered to S2 synapses (at time = 0) (n = 8). ***B*:** S2 TPS was delivered at the end of a 10-minute bath application of ISO (1.0 µM, indicated by the shaded region) (n = 9). ***C*:** S1 and S2 fEPSP slopes 45 minutes after S2 TPS from all experiments in *A* and *B*. A two-way ANOVA revealed a significant difference between S1 and S2 synapses (*F*_(1,30)_ = 38.962, *p* < 0.001), a significant effect of ISO (*F*_(1,30)_ = 33.212, *p* < 0.001), and a significant synapse x ISO interaction (*F*_(1,30)_ = 22.281, *p* < 0.001). ISO significantly enhanced LTP induction at S2 synapses (\*\**p* < 0.001) but had no lasting effect on S1 synapses (NS, not significant, *p* = 0.467, SNK *post hoc* tests). ***D*:** Control experiments where 15 seconds of TPS was delivered to S1 synapses prior to S2 TPS (n = 9). ***E*:** 15 seconds of S1 TPS before S2 TPS was delivered at the end of a 10-minute bath application of ISO (n = 10). ***F*:** S1 and S2 fEPSP slopes 45 minutes post-S2 TPS from all experiments in D and E. A two-way ANOVA with SNK *post hoc* tests revealed a significant difference between S1 and S2 synapses (*F*_(1,34)_ = 55.723, *p* < 0.001) and a significant synapse x ISO interaction (*F*_(1,34)_ = 62.341, *p* < 0.001). ISO significantly enhanced LTP induction at S1 synapses and inhibited LTP induction at S2 synapses (\*\**p* < 0.001). Histograms show number of EPSP evoking 0-4 dendritic PSs during S2 TPS. EPSP-evoked CS bursting during S2 TPS was significantly enhanced by prior S1 TPS (\**p* < 0.05, one-way ANOVA on Ranks with Dunns *post hoc* comparisons to control experiments shown in panel A, *H*_(3)_ = 30.874, *p* < 0.001). Traces show superimposed fEPSPs evoked by S1 and S2 stimulation during baseline and 45 minutes post-S2 TPS (calibration bars are 2 mV and 5 ms).

β-AR activation had two prominent effects on postsynaptic responses evoked during S1 and S2 TPS that likely contribute to the bidirectional effects of ISO on LTP induction shown in figure 3E and 3F. First, β-AR activation increased EPSP-evoked CS bursting during the 15 seconds of TPS delivered to S1 synapses (Fig. 4A). Given the crucial role of EPSP-evoked CS bursting in TPS-induced LTP (Thomas et al., 1998; O’Dell, 2022), the increase in CS bursting induced by β-AR activation likely contributes to the ISO-induced facilitation of LTP induction at S1 synapses, as suggested by others (Winder et al., 1999; Gelinas et al., 2008). Second, 15 seconds of S1 TPS in the presence of ISO induced a strong, heterosynaptic depression of transmission at S2 synapses that was maintained throughout the 5-seconds-long train of TPS delivered to S2 synapses (Fig. 4B). Interestingly, there was a significant, negative correlation between the potentiation of S2 synapses and the amount of heterosynaptic depression induced by S1 TPS (Fig. 4C). Moreover, the amount of heterosynaptic depression at S2 synapses was highly correlated with the probability of EPSP-evoked CS bursting during S1 TPS (Fig. 4D). Together, these correlations suggest that β-AR activation facilitates synaptic competition and suppresses LTP induction at S2 synapses by enhancing a CS burst-dependent form of heterosynaptic depression.

**Figure 4.**
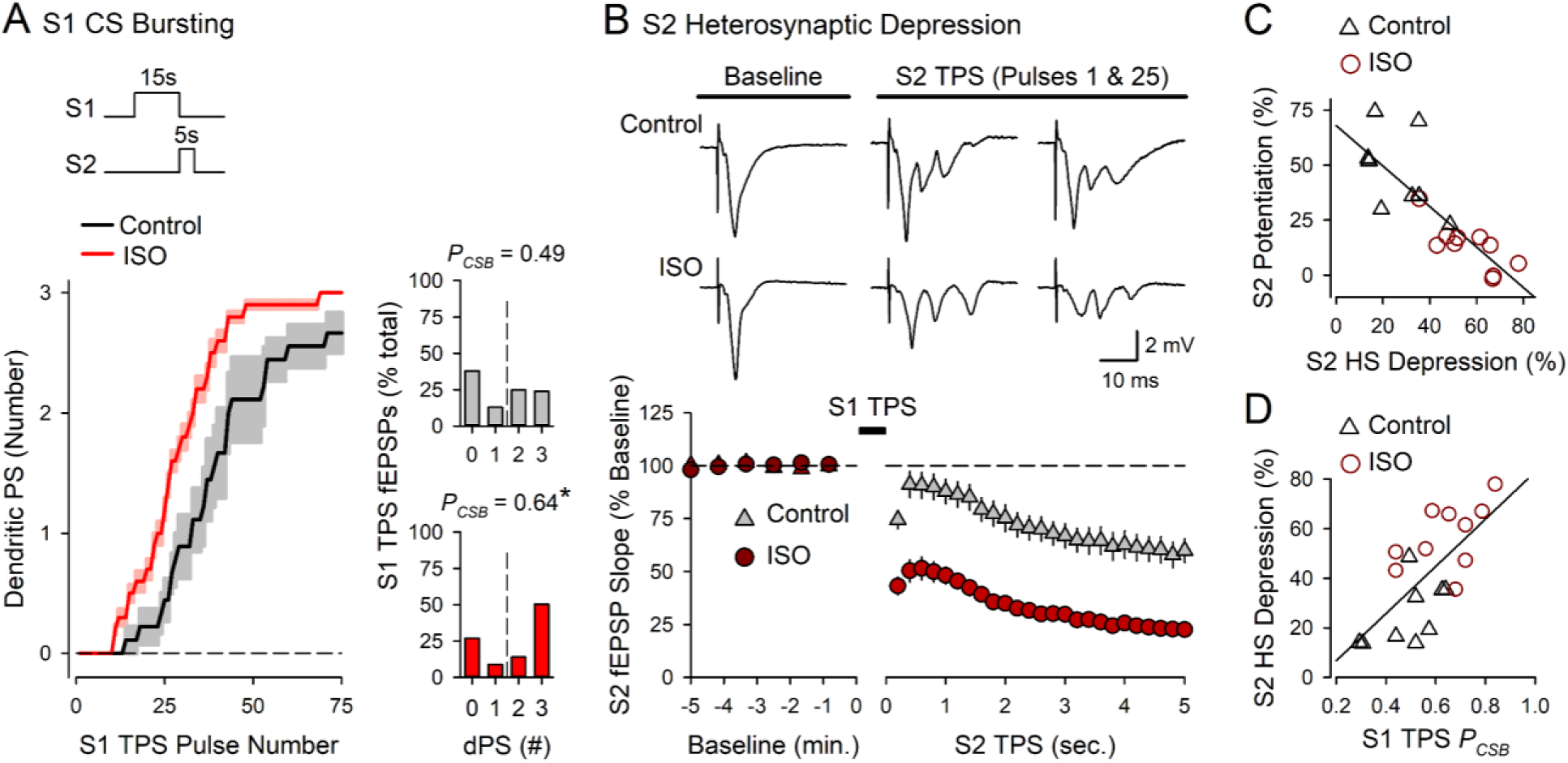
β-AR activation enhances EPSP-evoked CS bursting and TPS-induced heterosynaptic depression. Results are from the experiments shown in figure 3D and 3E. ***A*:** Number of dendritic PSs evoked by EPSPs during 15 s of TPS delivered to S1 synapses in the absence (control) and presence of 1.0 µM ISO. Shading represents ± SEM. Histograms show the number of EPSPs eliciting 0 – 3 dendritic PSs during S1 TPS in control experiments (*P*_CSB_ = 0.490) and S1 TPS in the presence ISO (*P*_CSB_ = 0.643, *t*_(17)_ = 2.538, \**p* = 0.0212). ***B*:** fEPSP slopes during S2 TPS delivered after 15 seconds of S1 TPS in the absence and presence of ISO. Traces show S2 fEPSPs evoked before S1 TPS (baseline) and the first and last fEPSPs evoked during S2 TPS. ***C* and *D*:** Scatter plots show correlations between S1 TPS-induced heterosynaptic (HS) depression and LTP induction at S2 synapses (*C*) (*r* = -0.837, *p* = 7.75 x 10^-6^) and between the probability of EPSP-evoked CS bursting during S1 TPS and the magnitude of heterosynaptic depression at S2 synapses (*D*) (*r* = 0.696, *p* = 9.4 x 10^-4^).

### 3.3 β-AR activation enhances a CS burst-dependent form of heterosynaptic depression

To directly test the notion that heterosynaptic depression is induced by postsynaptic CS bursting I examined whether suppressing EPSP-evoked CS bursting with a low concentration of the Na^+^ channel blocker TTX (Thomas et al., 1998; O’Dell, 2022) inhibits TPS-induced heterosynaptic depression. In these experiments S2 synapses were activated at 0.033 Hz before and after a 15-seconds-long train of TPS delivered to S1 synapses. In control experiments, 15 seconds of S1 TPS delivered at the end of a 10-minute bath application of 1.0 µM ISO induced strong EPSP-evoked bursting (*P_CSB_* = 0.66, n = 9) and a large, transient depression at S2 synapses (S2 fEPSPs were initially depressed to 45 ± 5% of baseline and recovered with a time constant of approximately 46 seconds) (Fig. 5A,B). Consistent with the notion that β-AR activation enhances a CS burst-dependent form of heterosynaptic depression, CS bursting during S1 TPS was blocked (*P_CSB_* during S1 TPS = 0.07, n = 6) and the heterosynaptic depression of S2 synapses was abolished when ISO was co-applied with 200 nM TTX (the first S2 fEPSPs evoked after S1 TPS were 98 ± 2% of baseline) (Fig. 5A,B). Notably, suppressing feedforward, GABAergic inhibition enhances EPSP-evoked CS bursting in CA1 pyramidal cells (Lovett-Barron et al., 2012; Babiec et al., 2017). Thus, as an additional test of the role of postsynaptic CS bursting in heterosynaptic depression I examined whether heterosynaptic depression is enhanced when S1 TPS is delivered in the presence of the GABA_A_ receptor antagonist gabazine. In control experiments, 15 seconds of S1 TPS induced a relatively modest amount of EPSP-evoked CS bursting (*P_CSB_* = 0.51, n = 9) and a small heterosynaptic depression of transmission at S2 synapses (S2 fEPSPs were initially depressed to 76 ± 2% of baseline) (Fig. 5C,D). In contrast, EPSP-evoked CS bursting was significantly enhanced when S1 TPS was delivered at the end of a 10-minute bath application of 0.5 µM gabazine (*P_CSB_* = 0.93) and, consistent with the notion that TPS induces a CS burst-dependent form of heterosynaptic depression, the first S2 fEPSPs evoked post-S1 TPS were reduced to just 33 ± 4% of baseline (n = 9) (Fig. 5C,D).

**Figure 5.**
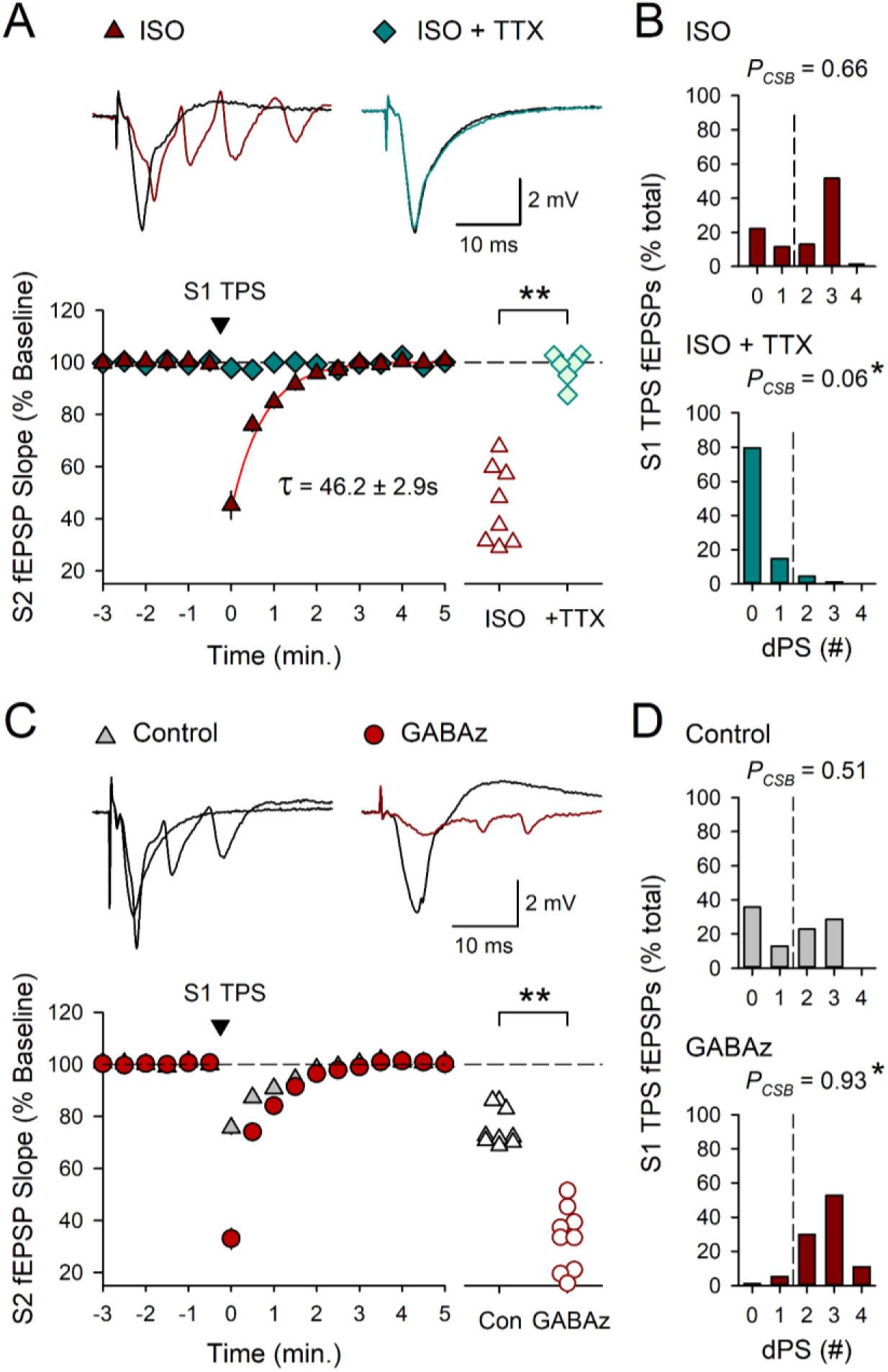
β-AR activation enhances a CS burst-dependent form of heterosynaptic depression. S2 synapses were activated at 0.033 Hz before and after a 15 s train of S1 TPS. ***A*:** S1 TPS delivered at the end of a 10-minute bath application of 1.0 µM ISO (n = 8) or ISO plus 0.2 µM TTX (n = 6). Red line shows a single exponential fit to the recovery of S2 responses following S1 TPS-induced heterosynaptic depression. Scatter plot shows slopes of first S2 fEPSPs evoked post-S1 TPS from all experiments (*t*_(12)_ = 8.051, \*\**p* = 3.52 x 10^-6^). ***B*:** Number of EPSPs eliciting 0 – 4 dendritic PSs during S1 TPS in the presence of ISO (*P*_CSB_ = 0.662) and ISO + TTX (*P*_CSB_ = 0.056, Mann-Whitney *U* = 0.0, \**p* < 0.001). ***C*:** S1 TPS was delivered either alone (control, n = 9) or at the end of a 10-minute bath application of 0.5 µM gabazine (GABAz) (n = 9). Scatter plot shows slopes of first S2 fEPSPs evoked post-S1 TPS from all experiments (*t*_(16)_ = 9.098, \*\**p* = 1.01 x 10^-7^). ***D*:** Number of EPSPS eliciting 0 – 4 dendritic PSs during S1 TPS in control experiments (*P*_CSB_ = 0.514) and in the presence of gabazine (*P*_CSB_ = 0.934, Mann Whitney *U* = 2.5, \**p* < 0.001). Traces in *A* and *C* show superimposed S2 fEPSPs recorded before and after S1 TPS.

GABAergic inhibitory synaptic transmission opposes the induction of Hebbian LTP (Wigström and Gustafsson, 1984) and GABA_A_ receptor antagonists are thus often used to facilitate LTP induction in in-vitro studies of synaptic plasticity, including BTSP (Bittner et al., 2017; Xiao et al., 2023). The hypothesis that heterosynaptic depression underlies synaptic competition predicts, however, that the robust increase in TPS-induced heterosynaptic depression induced by gabazine should have the opposite effect and inhibit, rather than enhance, the induction of LTP by cooperative synaptic interactions during TPS. To test this prediction, I examined the effects of gabazine on the heterosynaptic facilitation of LTP induction at S2 synapses produced by prior S1 TPS. As expected, in control experiments 15 seconds of S1 TPS delivered prior to 5 seconds of S2 TPS induced LTP at S1 synapses (S1 fEPSPs potentiated to 164 ± 6% of baseline, n = 7) and enabled LTP induction at S2 synapses (S2 fEPSPs potentiated to 152 ± 7% of baseline) (Fig. 6A). However, when this pattern of S1→S2 TPS was delivered at the end of a 10-minute bath application of gabazine the induction of LTP at S1 synapses was enhanced (S1 fEPSPs potentiated to 199 ± 6% of baseline, n = 6) and LTP induction at S2 synapses was abolished (45 minutes post-TPS S2 fEPSPs were 105 ± 4% of baseline) (Fig. 6B,C). Thus, consistent with the notion that synaptic competition is mediated by a CS burst-dependent form of heterosynaptic depression, the robust increase in S1 TPS-induced CS bursting and heterosynaptic depression induced by gabazine is associated with a strong increase in synaptic competition that inhibits LTP induction at S2 synapses.

**Figure 6.**
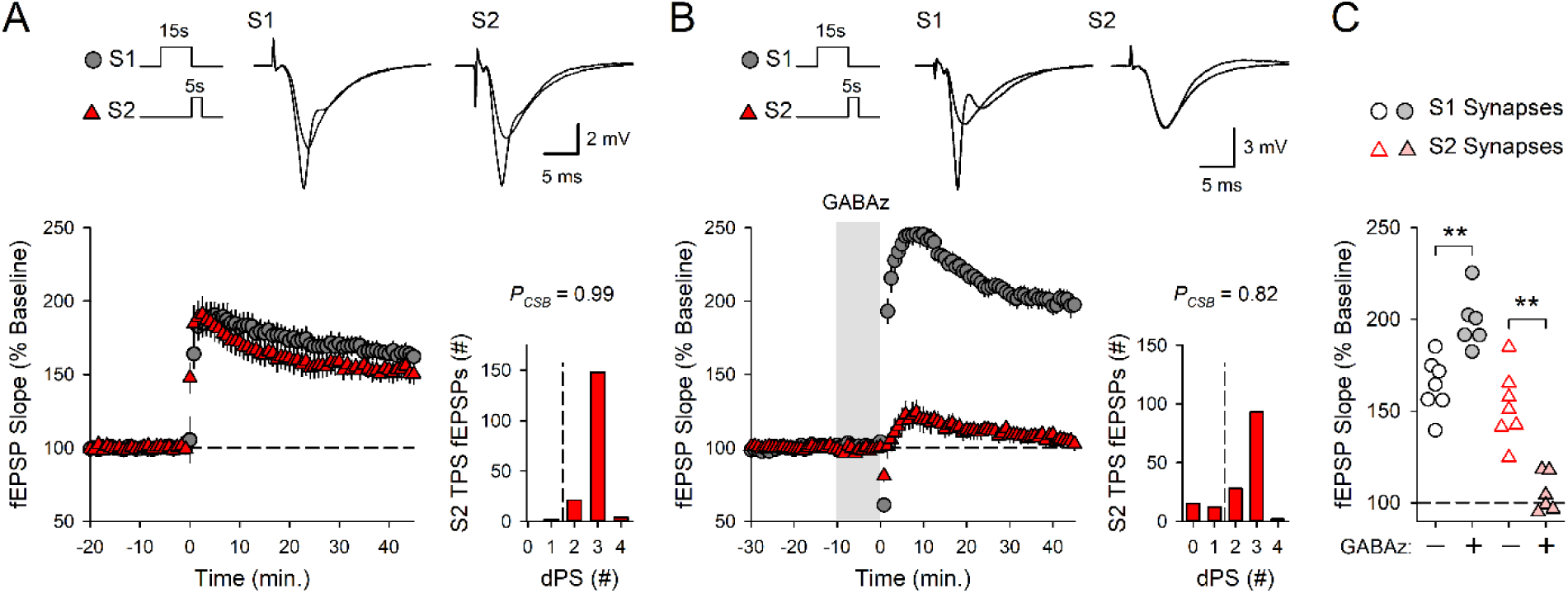
Suppressing GABAergic transmission enhances both LTP induction and synaptic competition. ***A*:** Control experiments (n = 7) where a 15-seconds-long train of TPS was delivered to S1 synapses prior to S2 TPS (at time = 0). ***B*:** 15 seconds of S1 TPS before S2 TPS was delivered at the end of a 10-minute bath application of 0.5 µM gabazine (GABAz) (n = 6). Histograms in *A* and *B* show number of EPSPs evoking 0-4 dendritic PSs during S2 TPS and traces show superimposed fEPSPs elicited by S1 and S2 stimulation during baseline and 45 minutes post-S2 TPS. ***C*:** S1 and S2 fEPSPs 45 minutes after S2 TPS from all experiments. A two-way ANOVA with SNK *post hoc* tests revealed a significant difference between S1 and S2 synapses (*F*_(1,22)_ = 74.892, *p* < 0.001) and a significant synapse x gabazine interaction (*F*_(1,22)_ = 45.834, *p* < 0.001). Gabazine significantly enhanced LTP at S1 synapses and suppressed LTP induction at S2 synapses (\*\**p* < 0.001).

### 3.4 Synaptic competition is mediated by a β-AR-modulated and adenosine receptor-dependent form of heterosynaptic depression

How might a CS burst-dependent form of heterosynaptic depression give rise to competitive synaptic interactions during the induction of CS burst-dependent LTP? One prominent form of heterosynaptic depression at excitatory synapses onto CA1 pyramidal cells is due to an inhibition of glutamate release caused by activation of presynaptic, A1-type adenosine receptors (Grover and Teyler, 1993; Mitchell et al., 1993; Wu and Saggau, 1994). Moreover, bursts of postsynaptic action potentials trigger adenosine release from CA1 pyramidal cell dendrites (Lovatt et al., 2012; Wu et al., 2023). Thus, synaptic competition induced by S1 TPS in the presence of ISO may be mediated by a presynaptic, adenosine receptor-dependent form of heterosynaptic depression that, by inhibiting glutamate release, prevents the strong activation of postsynaptic NMDARs needed for LTP induction at S2 synapses. To test this hypothesis, I first examined the effects of the A1 adenosine receptor antagonist DPCPX on the heterosynaptic depression induced by TPS. Consistent with the notion that TPS-induced heterosynaptic depression in due to activation of A1 receptors, the heterosynaptic depression induced by 15 seconds of S1 TPS in the presence of ISO (S2 fEPSPs were initially reduced to 42 ± 5% of baseline, n = 7) was significantly inhibited when S1 TPS was delivered in the presence of ISO plus 400 nM DPCPX (the first S2 fEPSPs elicited post-S1 TPS were 92 ± 4% of baseline, n = 7) (Fig. 7A). To explore the role of heterosynaptic depression in synaptic competition I next examined the effects of DPCPX on LTP induction when 15 seconds of S1 TPS was delivered before 5 seconds of S2 TPS in the presence of ISO. In control experiments, 15 seconds of S1 TPS induced LTP at S1 synapses (fEPSPs potentiated to 202 ± 5% of baseline, n = 7) and suppressed LTP induction at S2 synapses (45 minutes post-TPS, S2 fEPSPs were just 110 ± 2% of baseline) (Fig. 7B). In contrast, when this same pattern of S1 → S2 TPS was delivered in the presence of ISO plus DPCPX, S1 synapses potentiated to 194 ± 11% of baseline and S2 synapses potentiated to 194 ± 8% of baseline (n = 8) (Fig. 7C). Thus, although DPCPX had no effect on LTP induction at S1 synapses, it blocked the enhancement of synaptic competition induced by β-AR activation, thereby enabling LTP induction at S2 synapses (Fig. 7D). Importantly, DPCPX did not enable the induction of LTP when 5 seconds of TPS was delivered to S2 synapses without prior S1 TPS (Fig. 7E). This indicates that blocking A1 receptors does not simply enhance LTP induction but instead selectively disrupts the increase in synaptic competition induced by β-AR activation.

**Figure 7.**
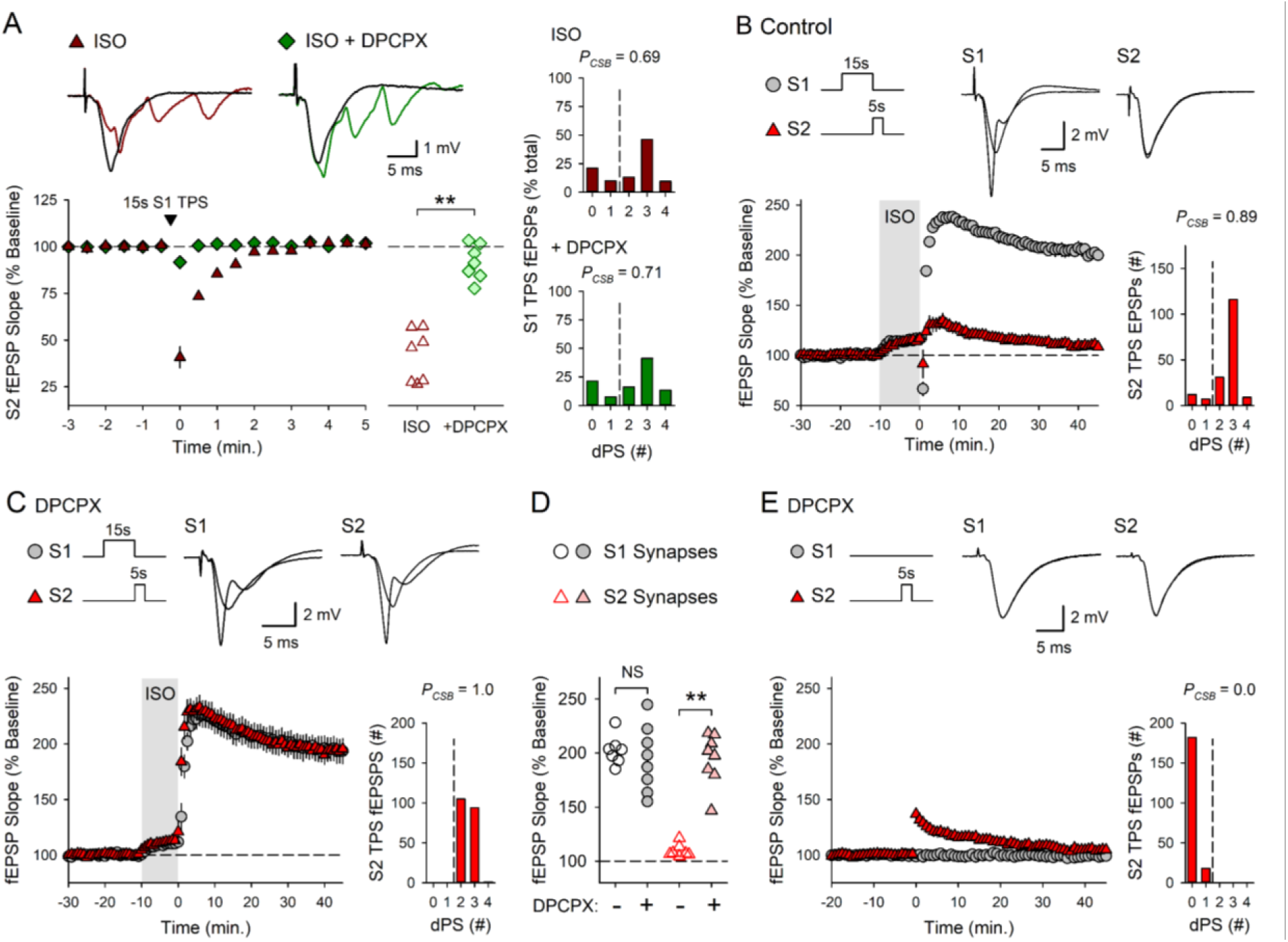
Synaptic competition is mediated by an A1 adenosine receptor-dependent form of heterosynaptic depression. ***A*:** S2 synapses were activated at 0.033 Hz before and after a 15-seconds-long train of S1 TPS (delivered at time = 0 after a 10-minute bath application of 1.0 µM ISO). The heterosynaptic depression induced by S1 TPS in control experiments (ISO alone, n = 7) was inhibited when S1 TPS was delivered in the presence of ISO plus 400 nM DPCPX (n = 7). Scatter plot shows slopes of first S2 fEPSPs evoked post-S1 TPS from all experiments (*t*_(12)_ = 7.875, \*\**p* = 4.42×10^-6^). Histograms show the number of EPSPs eliciting 0 – 4 dendritic PSs during S1 TPS in the presence of ISO (*P*_CSB_ = 0.69) and in the presence ISO plus DPCPX (*P*_CSB_ = 0.71, *t*_(12)_ = 0.287, *p* = 0.779). Traces show superimposed S2 fEPSPs recorded during baseline and after S1 TPS. ***B*:** Control experiments where 15 seconds of TPS was delivered to S1 synapses prior to 5 seconds of S2 TPS at the end of a 10-minute bath application of 1.0 µM ISO (n = 7). ***C*:** 15 seconds of S1 TPS before 5 seconds of S2 was delivered at the end of a 10-minute bath application of 1.0 µM ISO in slices continuously bathed in ACSF containing 400 nM DPCPX (n = 8). Traces in B and C show superimposed fEPSPs elicited by S1 and S2 stimulation during baseline and 45 minutes post-S2 TPS. Histograms in B and C show the number of EPSPs evoking 0-4 dendritic PSs during S2 TPS. ***D*:** Summary plot showing fEPSP slopes recorded 45 minutes post-S2 TPS from all experiments in B and C. A two-way ANOVA with SNK *post hoc* tests revealed a significant difference between S1 and S2 synapses (*F*_(1,26)_ = 35.706, *p* < 0.001), a significant effect of DPCPX (*F*_(1,26)_ = 24.628, *p* < 0.001), and a significant synapse x DPCPX interaction (*F*_(1,26)_ = 35.864, *p* < 0.001). DPCPX significantly enhanced LTP induction at S2 synapses (**p < 0.001) but had no effect on LTP induction at S1 synapses (NS, *p* = 0.475). **E:** 5 seconds of S2 TPS was delivered alone (at time = 0) in slices bathed in ACSF containing 400 nM DPCPX (n = 8). 45 minutes post-TPS S2 fEPSPs were 105 ± 2% of baseline. Traces show superimposed fEPSPs elicited by S1 and S2 stimulation during baseline and 45 minutes post-TPS.

### 3.5 β-AR activation enhances cooperative synaptic interactions during the induction of CS burst-dependent LTP

Although delivering 15 seconds of TPS to S1 synapses before 5 seconds of S2 TPS induces similar amounts of LTP at both synapses in control experiments (Fig. 3D and 6A), the enhancement of synaptic competition induced by β-AR activation results in a selective potentiation of the more strongly activated, S1 synapses (Fig. 3E and 7B). This differential potentiation of more strongly activated synapses when TPS is delivered in the presence of ISO raises an intriguing question – how does β-AR activation regulate synaptic interactions and LTP induction when S1 and S2 synapses are activated by more similar, or even identical, trains of TPS? To address this question, I examined how β-AR activation influences LTP induction at S1 and S2 synapses when the duration of TPS delivered to S1 synapses (before 5 seconds of S2 TPS) was reduced to 10 seconds. Consistent with the results shown in figure 2B, in control experiments 10 seconds of S1 TPS induced a modest potentiation at S1 synapses (S1 fEPSPs potentiated to 129 ± 3% of baseline, n = 8) and enabled LTP induction at S2 synapses (S2 fEPSPs potentiated to 164 ± 6% of baseline (Fig. 8A). However, when this pattern of S1→S2 TPS was delivered in the presence of ISO, β-AR activation significantly enhanced LTP induction at S1 synapses (S1 fEPSPs potentiated to 169 ± 7% of baseline, n = 10) but had no effect on LTP induction at S2 synapses (S2 fEPSPs potentiated to 172 ± 6% of baseline) (Fig. 8B,C). The enhancement of synaptic competition induced by β-AR activation thus appears to be highly activity-dependent. Indeed, β-AR activation significantly enhanced LTP induction at both S1 and S2 synapses when the duration of S1 TPS was further reduced to 5 seconds (Fig. 8D-F). In control experiments, 5 seconds of S1 TPS delivered before 5 seconds of S2 TPS had no lasting effect on synaptic transmission at S1 synapses and did not enable LTP induction at S2 synapses (45 minutes post-TPS S1 and S2 synapses were 107 ± 4% and 113 ± 4% of baseline, respectively, n = 10) (Fig. 8D). In the presence of ISO, this pattern of S1 → S2 TPS induced LTP at both S1 and S2 synapses and, surprisingly, the potentiation at S2 synapses was significantly larger than that seen as S1 synapses (S1 and S2 synapses potentiated to 161 ± 5% and 189 ± 6% of baseline, respectively, n = 10) (Fig. 8E,F). Moreover, although 5 seconds of S1 TPS induced a modest heterosynaptic facilitation of EPSP-evoked CS bursting at S2 synapses in control experiments (*P_CSB_* = 0.29) (Fig. 8D), EPSP-evoked CS bursting during S2 TPS was enhanced when S1 and S2 TPS trains were delivered in the presence of ISO (*P_CSB_* = 0.65), although this effect was marginally significant (*p* = 0.049) (Fig. 8E). Together, these results indicate that β-AR activation enhances cooperative synaptic interactions during brief trains of TPS that are normally below threshold for LTP induction. Consistent with this, the potentiation of S2 synapses induced by S1→ S2 TPS in the presence of ISO (Fig. 8E) was also significantly larger than that induced by 5 seconds of S2 TPS delivered in the presence of ISO without prior S1 TPS (Fig. 3B) (*t*_(17)_ = 3.606, *p* = 0.00218).

**Figure 8.**
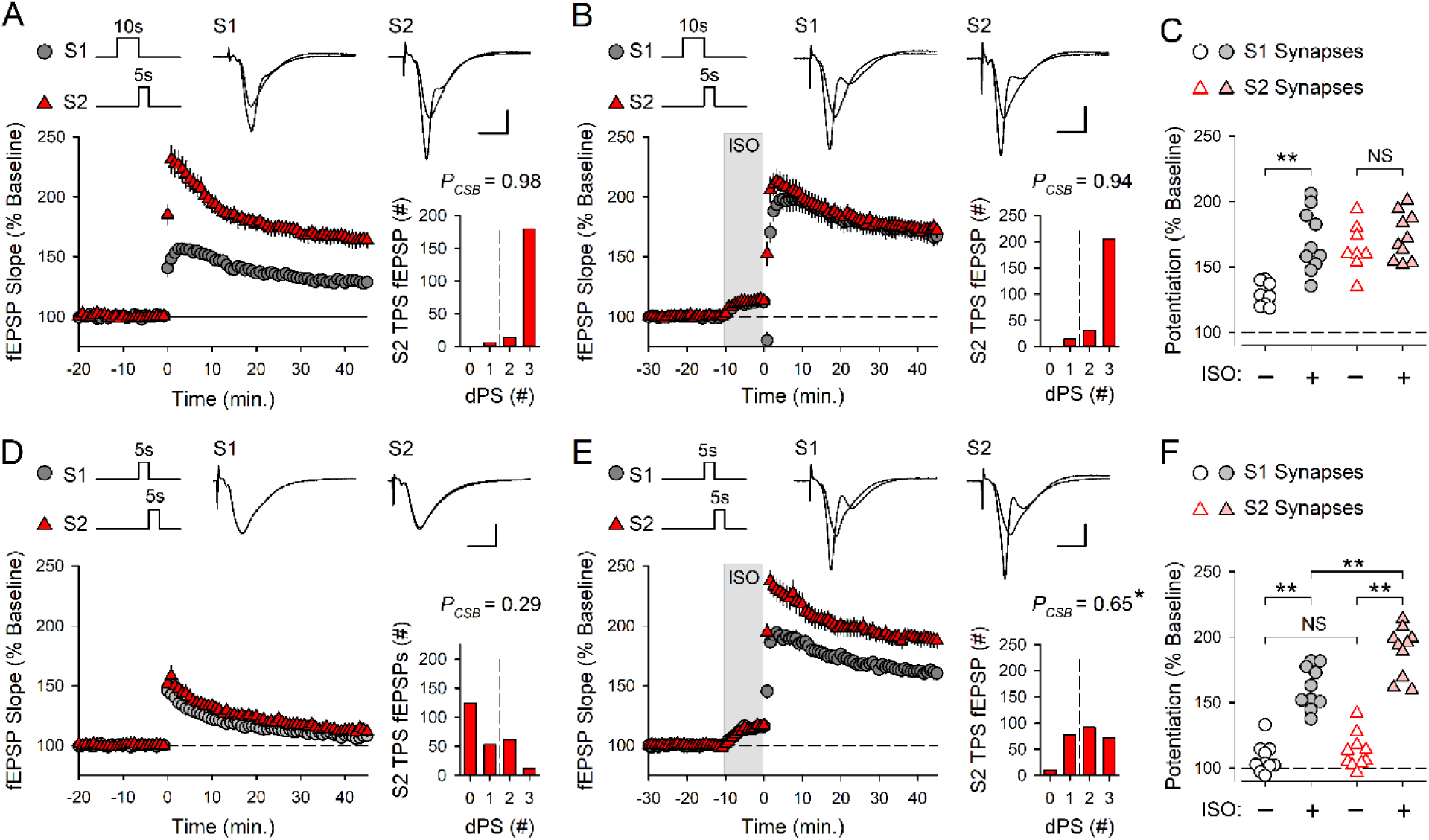
β-AR activation induces an activity-dependent facilitation of cooperative synaptic interactions during CS burst-dependent LTP induction. ***A*:** Control experiments (n = 8) where 10 seconds of TPS was delivered to S1 synapses before 5 seconds of S2 TPS (at time = 0). ***B*:** 10 seconds of S1 TPS before S2 TPS was delivered at the end of a 10-minute bath application of ISO (n = 10). ***C*:** S1 and S2 fEPSP slopes 45 minutes after S2 TPS from all experiments in *A* and *B*. Results were analyzed a using two-way ANOVA with SNK *post hoc* comparisons (\*\**p* < 0.001, NS, not significant, *p* = 0.679). There was a significant difference between S1 and S2 synapses (*F*_(1,32)_ = 9.906, *p* = 0.004), a significant effect of ISO (*F*_(1,32)_ = 15.516, *p* < 0.001), and a significant synapse x ISO interaction (*F*_(1,32)_ = 6.712, *p* = 0.014). ***D*:** Control experiments where 5 seconds of S1 TPS was delivered before 5 seconds of S2 TPS (n = 10). ***E*:** 5 seconds of S1 TPS before S2 TPS was delivered in the presence of ISO (n = 10). **F:** S1 and S2 fEPSP slopes 45 minutes after S2 TPS from all experiments in *D* and *E*. A two-way ANOVA with SNK post hoc comparisons (\*\**p* < 0.001, NS, not significant, *p* = 0.439) revealed a significant difference between S1 and S2 synapses (*F*_(1,36)_ = 11.626, *p* = 0.002), a significant effect of ISO (*F*_(1,36)_ = 181.462, *p* < 0.001, and a significant synapse x ISO interaction (*F*_(1,36)_ = 5.307, *p* = 0.027). Histograms show number of EPSPs evoking 0 – 3 CS bursts during S2 TPS in all experiments. CS bursting during S2 TPS delivered after of 5 seconds of S1 TPS was significantly enhanced by ISO (Mann Whitney *U* = 24.0, \**p* = 0.049). Traces show superimposed fEPSPs elicited by S1 and S2 stimulation during baseline and 45 minutes after S2 TPS.

As an additional test of the ability of β-AR activation to enhance cooperativity in CS burst-dependent LTP, I also examined whether ISO regulates cooperative synaptic interactions using a very brief train of S2 TPS (2 seconds) that, by itself, fails to induce LTP, even when delivered in the presence of ISO (Fig. 9). In control experiments, 2 seconds of S2 TPS alone failed to elicit CS bursts (*P_CSB_* = 0.0, n = 9) and had no lasting effect on synaptic strength (45 minutes post-TPS S2 fEPSPs were 105 ± 3% of baseline) (Fig. 9A). As expected, 10 seconds of S1 TPS delivered before S2 TPS facilitated EPSP-evoked CS bursting during S2 TPS (*P_CSB_* = 0.85, n = 8) and enabled the induction of a modest, but significant, potentiation of S2 synapses (S2 fEPSPs potentiated to 127 ± 4% of baseline, *p* < 0.001 compared to S2 TPS alone) (Fig. 9B). In contrast, β-AR activation did not enable LTP induction when 2 seconds of S2 TPS was delivered without prior S1 TPS (45 minutes post-TPS S2 fEPSPs were 104 ± 3% of baseline, n = 7) (Fig. 9C). β-AR activation did, however, significantly enhance the heterosynaptic facilitation of LTP induction at S2 synapses induced by 10 seconds of S1 TPS (S2 fEPSPs potentiated to 166 ± 4% of baseline, n = 9) (Fig. 9D,E). Together with the results shown in figure 8, the highly synergistic effects of ISO and S1 TPS on LTP induction at S2 synapses indicate that β-AR activation strongly enhances cooperative synaptic interactions during the induction of CS burst-dependent LTP. Indeed, the combined effects of ISO and S1 TPS enables the induction of a robust (> 60% increase) and persistent potentiation at S2 synapses by TPS trains containing just 10 pulses of presynaptic fiber stimulation (Fig. 9D).

**Figure 9.**
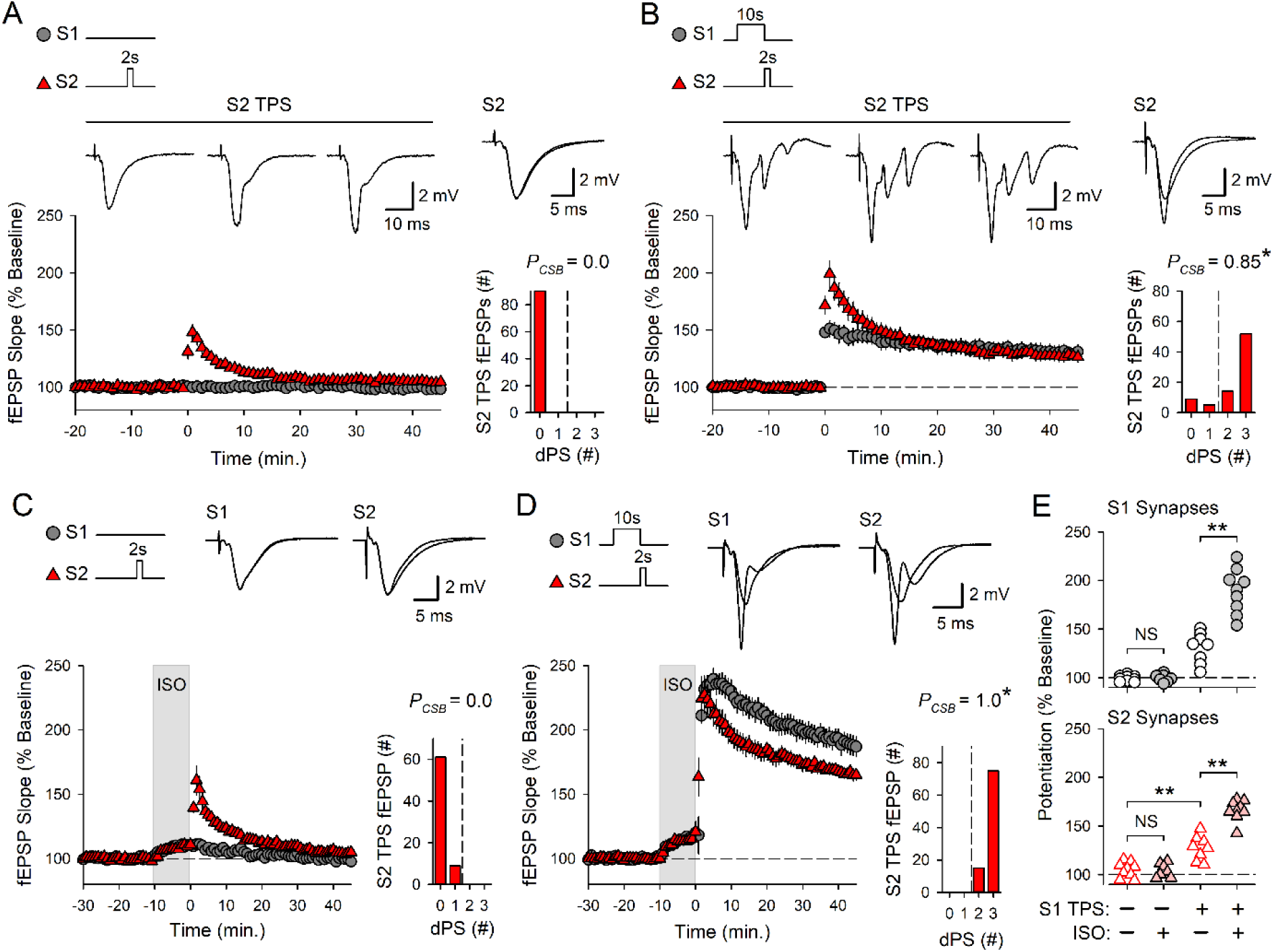
β-AR activation and TPS interact in a highly synergistic fashion to promote LTP induction by cooperative synaptic interactions. ***A*:** Control experiments (n = 9) where 2 seconds of TPS was delivered to S2 synapses (at time = 0). ***B*:** 10 seconds of TPS was delivered to S1 synapses before 2 seconds of S2 TPS (n = 8). Traces in A and B show fEPSPs evoked by the 1^st^, 5^th^, and 10^th^ stimulation pulses during S2 TPS and superimposed S2 fEPSPs evoked during baseline and 45 minutes after S2 TPS. ***C*:** 2 seconds of TPS was delivered to S2 synapses at the end of a 10-minute bath application of ISO (n = 7). ***D*:** 10 seconds of S1 TPS before 2 seconds of S2 TPS was delivered at the end of a 10-minute bath application of ISO (n = 9). Traces in *C* and *D* show superimposed fEPSPs elicited by S1 and S2 stimulation during baseline and 45 minutes post-S2 TPS. Histograms in *A*-*D* show number of EPSPs evoking 0 – 3 dendritic PSs during S2 TPS. \**p* < 0.05 compared to control (A), one-way ANOVA on Ranks, *H*_(3)_ = 28.182, p < 0.001. ***E*:** S1 (top) and S2 (bottom) fEPSP slopes recorded 45 minutes after S2 TPS from all experiments. Results for S1 and S2 synapses were analyzed separately using two-way ANOVAs followed by SNK comparisons (\*\**p* < 0.001, NS, not significant, *p* = 0.973 for S1 synapses and *p* = 0.915 for S2 synapses). For S1 synapses there was a significant effect of S1 TPS (*F*_(1,29)_ = 144.244, *p* < 0.001), ISO (*F*_(1,29)_ = 33.750, *p* < 0.001), and a significant S1 TPS x ISO interaction (*F*_(1,29)_ = 33.188, *p* < 0.001). For S2 synapses there was a significant effect of S1 TPS (*F*_(1,29)_ = 149.566, *p* < 0.001), ISO (*F*_(1,29)_ = 30.693, *p* < 0.001), and a significant S1 TPS x ISO interaction (*F*_(1,29)_ = 32.442, *p* < 0.001).

### 3.6 β-AR activation prolongs the time course of cooperative synaptic interactions during the induction of CS burst-dependent LTP

β-AR activation not only enhances the induction of conventional Hebbian LTP but also increases the time window during which synapses can interact in a cooperative/associative fashion and undergo LTP (Lin et al., 2003). Although the temporal constraints on the ability of synapses to interact in a cooperative fashion to induce conventional Hebbian LTP and CS burst-dependent LTP are very different (tens of milliseconds vs. several seconds, respectively) (Lin et al., 2003, O’Dell, 2022), this raises the interesting possibility that β-AR activation also regulates the temporal properties of cooperativity in CS burst-dependent LTP. Importantly, the heterosynaptic facilitation of EPSP-evoked CS bursting induced by TPS underlies cooperative synaptic interactions during the induction of CS burst-dependent LTP (O’Dell, 2022). Thus, to determine whether β-AR activation regulates the temporal properties of cooperativity in CS burst-dependent LTP I first examined the effects of ISO on the heterosynaptic facilitation of EPSP-evoked CS bursting induced by TPS. In these experiments 10 seconds of TPS was delivered to S1 synapses to induce EPSP-evoked CS bursting and 2 seconds of TPS was then delivered to S2 synapses with inter-train intervals (ITIs) ranging from 2 – 15 seconds. In control experiments EPSP-evoked CS bursting during S2 TPS was strongly upregulated when S2 TPS was delivered 2-6 seconds after S1 TPS (Fig. 10A,B) but no longer significantly different from control (S2 TPS alone) when the S1-S2 TPS ITI was increased to 8 seconds (Fig. 10C). In contrast, when these same patterns of S1 and S2 TPS were delivered at the end of a 10-minute bath application of ISO, the heterosynaptic facilitation of EPSP-evoked CS bursting during S2 TPS was still robust when the S1-S2 TPS ITI was increased to 8 seconds (Fig. 10D-F) and only began to fade when the ITI was increased to 12 seconds or more (Fig. 10G). As shown in figure 11, the heterosynaptic facilitation of LTP induction at S2 synapses induced by S1 TPS in the presence ISO exhibited a similar time course, with S2 fEPSPs potentiating to 150% of baseline or more when S2 TPS was delivered 2-10 seconds after S1 TPS in the presence of ISO (Fig. 11A,B). However, LTP induction at S2 synapses was reduced when the S1-S2 TPS ITI was increased to 12 seconds and no longer significant from control (S2 TPS alone in the presence of ISO alone) when the S1-S2 TPS ITI was increased to 15 seconds (Fig. 11C,D). Consistent with the notion that LTP induction at S2 synapses is enabled by the heterosynaptic facilitation of EPSP-evoked CS bursting produced by S1 TPS, there was a highly significant correlation between the magnitude of LTP at S2 synapses and the probability of EPSP-evoked CS bursting during S2 TPS (Fig. 11E). Thus, β-AR activation not only prolongs the transient heterosynaptic facilitation of EPSP-evoked CS bursting induced by TPS (Fig. 10) but also produces a broad temporal window lasting at least 10 seconds during which independent synapses can interact with one another in a cooperative fashion to undergo CS burst-dependent LTP. Notably, the timescale of cooperativity in CS burst-dependent LTP in the presence of ISO is nearly 2-fold longer than that seen in control conditions (O’Dell, 2022) and approximately 3-orders of magnitude longer than the timescale of cooperativity in conventional Hebbian LTP (Lin et al., 2003).

**Figure 10.**
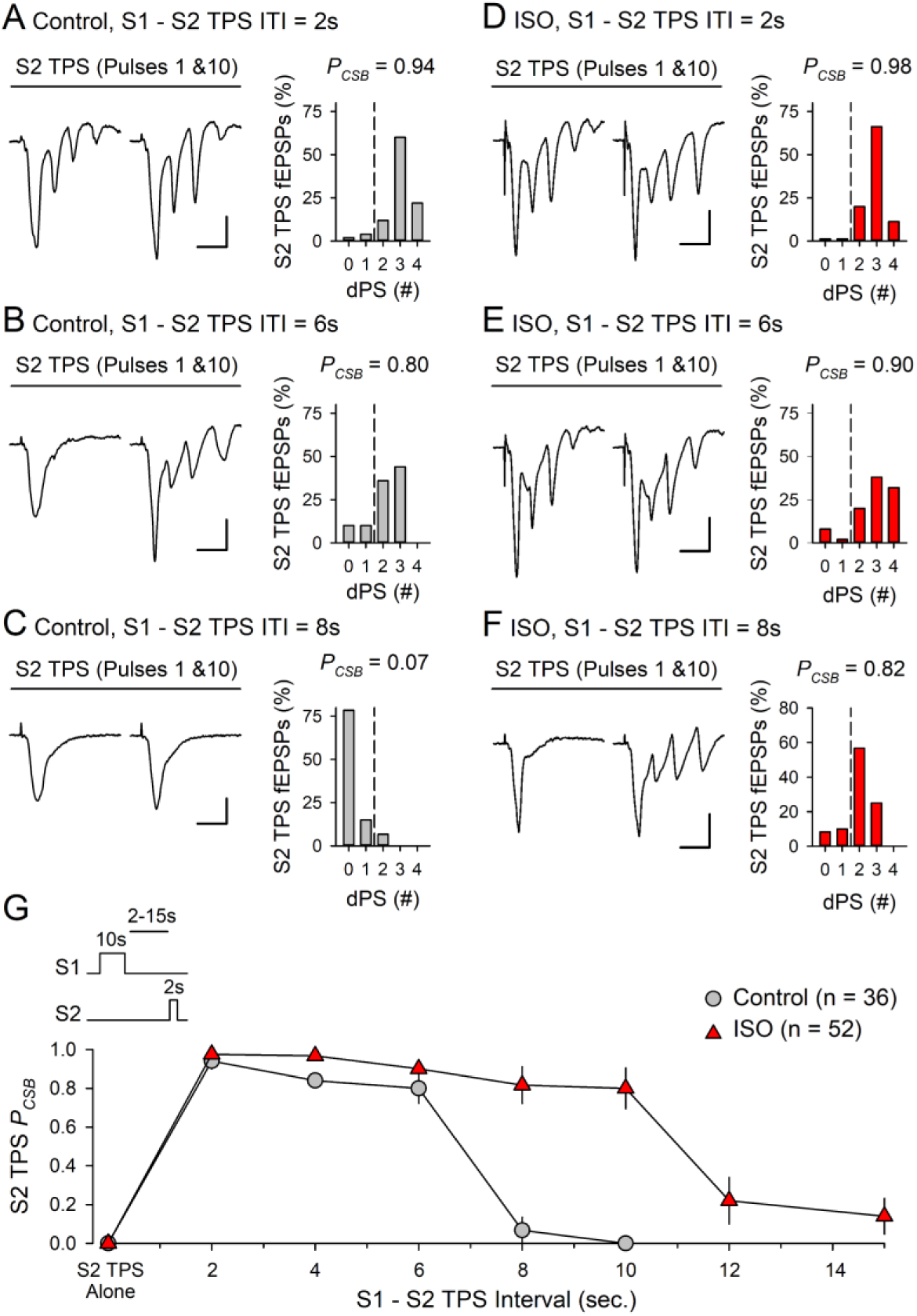
β-AR activation prolongs the short-term, heterosynaptic facilitation of EPSP-evoked CS bursting induced by TPS. ***A-C*:** Two seconds of S2 TPS was delivered 2 (A, n = 5), 6 (B, n = 5), or 8 seconds (C, n = 6) after a 10-seconds-long train of TPS delivered to S1 synapses. ***D-F*:** Results from experiments where the same patterns of S1→S2 TPS were delivered at the end of a 10-minute bath application of 1.0 µM ISO. N’s = 8, 5, and 6 for inter-train intervals (ITI) of 2, 6, and 8 seconds, respectively. Histograms in A-F show the number of EPSPs evoking 0-4 dendritic PSs during S2 TPS and traces show examples of the first and last fEPSPs evoked during S2 TPS. Calibration bars are 2 mV and 10 ms. ***G*:** Probability of CS bursting (*P_CSB_*) during S2 TPS trains delivered 2-15 seconds after S1 TPS in the presence and absence of ISO. In control experiments EPSP-evoked CS bursting during S2 TPS was significantly enhanced (*p* < 0.05, compared to S2 TPS alone results in Fig. 9A) with S1→S2 TPS ITIs up to 6 seconds (one-way ANOVA on Ranks with Dunn’s test comparisons to S2 TPS alone, *H*_(5)_ = 31.873, *p* < 0.001). In the presence of ISO, EPSP-evoked CS bursting during S2 TPS was significantly enhanced with S1→S2 ITIs up to 10 seconds (compared to S2 TPS alone in the presence of ISO, n = 9, *H*_(7)_ = 41.598, *p* < 0.001).

**Figure 11.**
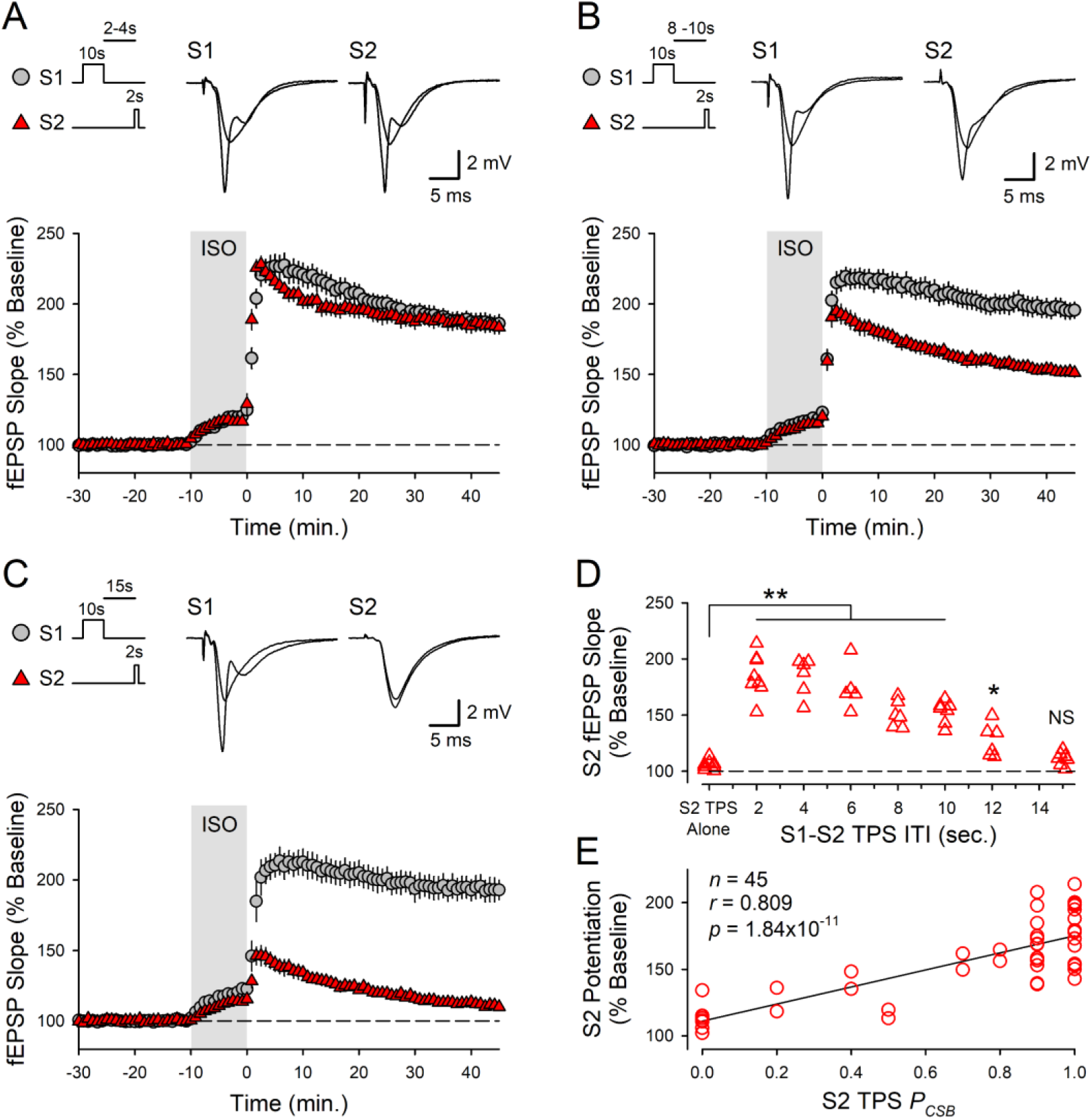
β-AR activation produces a broad time window for cooperative synaptic interactions during the induction CS burst-dependent LTP. ***A*:** Combined results from experiments where 2 seconds of S2 TPS was delivered 2 (n = 8) or 4 seconds (n = 6) after a 10-seconds-long train of S1 TPS in the presence of ISO. ***B*:** Same as in A but with S1→S2 inter-train intervals (ITIs) of 8 (n = 6) or 10 seconds (n = 7). ***C*:** S2 TPS (2 sec.) was delivered 15 seconds after 10 seconds of S1 TPS (n = 7). Traces in A – C show superimposed fEPSPs elicited by S1 and S2 stimulation during baseline and 45 minutes post-S2 TPS. ***D*:** S2 fEPSP slopes recorded 45 minutes post-S2 TPS from all experiments. Compared to control experiments (2 seconds of S2 TPS in the presence of ISO without prior S1 TPS, n = 7), S2 fEPSPs were significantly potentiated (\*\**p* < 0.001, \**p* = 0.041) with S1→S2 TPS ITIs up to 12 seconds (one-way ANOVA with SNK *post hoc* comparisons, *F*_(7,44)_ = 30.922, *p* < 0.001, NS, *p* = 0.545). ***E*:** Scatter plot shows the percent increase in synaptic strength at S2 synapses plotted as a function of *P*_CSB_ during S2 TPS for all experiments where S1 TPS trains were delivered before S2 TPS.

## 4 DISCUSSION

In agreement with earlier studies (Thomas et al., 1996; Winder et al., 1999; Gelinas and Nguyen, 2005; Gelinas et al., 2008; Qian et al., 2012; Jami et al., 2023), I find that β-AR activation enhances the induction of homosynaptic LTP by TPS protocols. However, the effects of ISO on synaptic interactions during different patterns of TPS indicate that β-AR activation exerts equally robust modulatory effects on activity-dependent forms of heterosynaptic plasticity that regulate CS burst-dependent LTP induction. Interestingly, although β-AR activation facilitated the induction of homosynaptic LTP at S1 synapses by trains of TPS lasting 5 or more seconds, it could enhance, suppress, or have no effect on the heterosynaptic facilitation of LTP induction at S2 synapses induced by S1 TPS. The effects of β-AR on synaptic interactions during CS burst-dependent LTP induction are thus surprisingly complex. A reanalysis of the effects of ISO on changes in synaptic strength induced by different patterns of S1→S2 TPS that includes results from experiments using trains of S1 TPS lasting 20 and 25 seconds (Supplemental Fig. 1) provides a useful context for thinking about these curious results (Fig. 12). In control experiments, the duration of TPS delivered to S1 synapses has markedly different effects on changes in synaptic strength at S1 and S2 synapses. At S1 synapses, LTP induction follows a simple rule – as the duration of S1 TPS is increased, the potentiation of S1 synapses grows (Fig. 12A). In contrast, brief trains of S1 TPS that induce modest LTP at S1 synapses induce a strong heterosynaptic facilitation of LTP induction at S2 synapses while longer trains of S1 TPS that induce more robust homosynaptic potentiation at S1 synapses inhibit LTP induction at S2 synapses. Thus, LTP induction at S2 synapses exhibits a pronounced, inverted-U shaped dependence on the duration of S1 TPS (Fig. 12B). In the presence of ISO, LTP induction at S1 synapses is enhanced and, as in control experiments, longer duration trains of S1 TPS tend to induce larger increases in synaptic strength (Fig. 12A). Notably, the inverted-U shaped relationship between S1 TPS duration and S2 synapse potentiation is also preserved when TPS is delivered in the presence of ISO (Fig. 12B). However, by enhancing both cooperative and competitive synaptic interactions, β-AR activation produces a pronounced leftward shift in the inverted-U shaped relationship between S1 TPS duration and S2 synapse potentiation (Fig. 12B).

**Figure 12.**
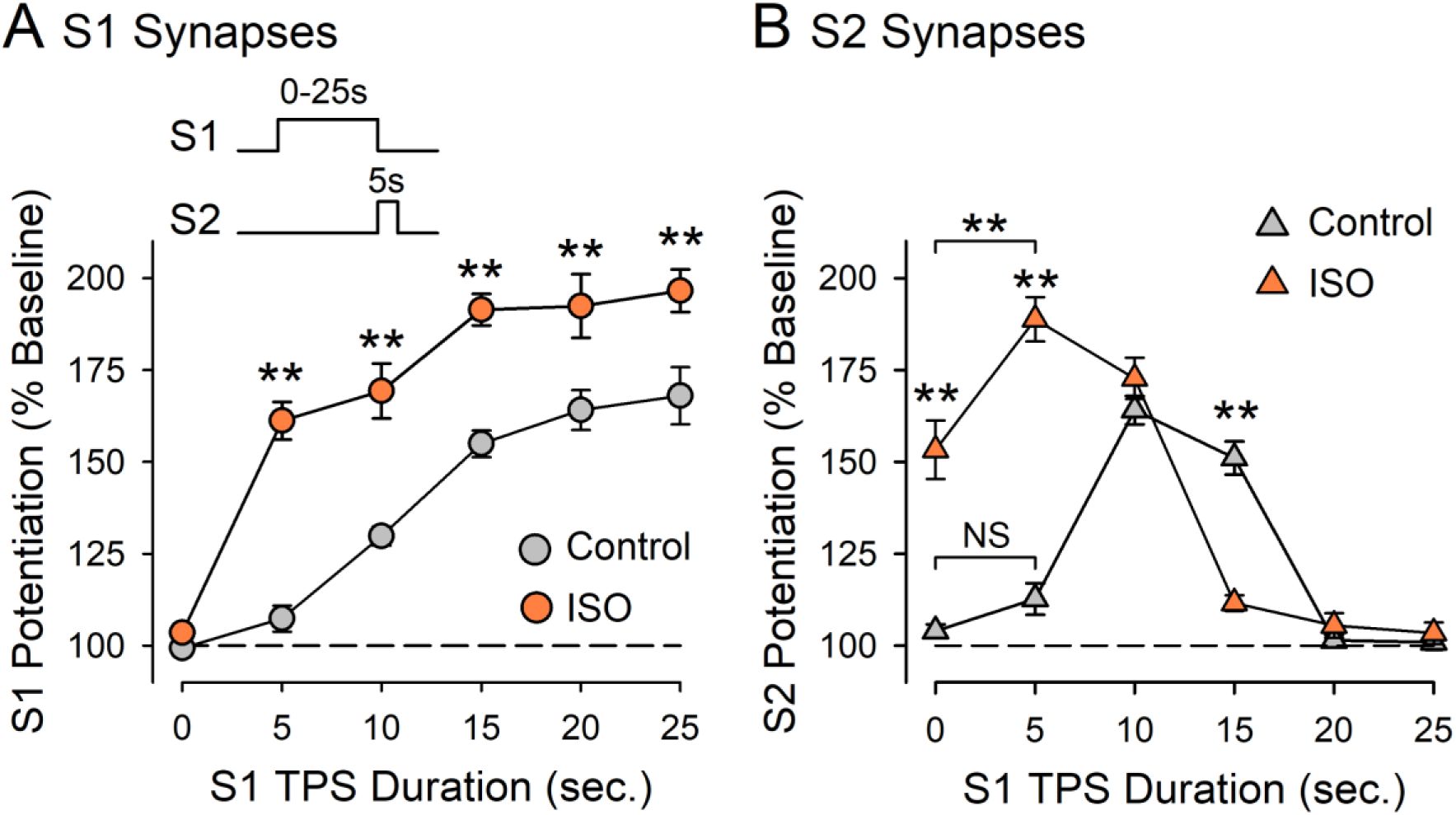
Effects of β-AR activation on synaptic interactions during CS burst dependent-LTP induction. Plots include results from experiments shown in figures 2, 3, 6A, 7B, 8, and supplemental figure 1 where different durations of S1 TPS were delivered before 5-seconds of S2 TPS. ***A*:** Changes in strength at S1 synapses induced by 0-25 seconds of S1 TPS delivered either alone (control) or in the presence of 1.0 µM ISO. A two-way ANOVA with SNK pairwise comparisons of control vs. ISO (\*\**p* < 0.001) indicated a significant effect of S1 TPS duration (*F*_(5,129)_ = 89.383, *p* < 0.001), a significant effect of ISO (*F*_(1,129)_ = 132.786, *p* < 0.001), and a significant S1 TPS duration x ISO interaction (*F*_(5,129)_ = 6.195, *p* < 0.001). ***B*:** Changes in S2 synaptic strength in the same experiments. A two-way ANOVA with SNK post-hoc comparisons revealed a significant effect of S1 duration (*F*_(5,129)_ = 73.289, *p* < 0.001), a significant effect of ISO (*F*_(1, 129)_ = 48.677, *p* < 0.001), and a significant S1 TPS duration x ISO interaction (*F*_(5,129)_ = 54.36, *p* < 0.001). 5 seconds of S1 TPS delivered before S2 TPS had no effect in control experiments (NS, *p* = 0.109) but facilitated LTP induction at S2 synapses when TPS was delivered in the presence of ISO (\*\**p* < 0.001). β-AR activation inhibited LTP induction at S2 synapses when S2 TPS followed 15 seconds of S1 TPS (\*\**p* < 0.001). *N* = 79 control experiments and 62 experiments where TPS protocols were delivered in the presence of ISO.

Interestingly, the analysis in figure 12 indicates that while β-AR activation enhances CS burst-dependent LTP, it but does so in a way that maintains the relationship between the magnitude of homosynaptic LTP and synaptic competition (i.e., synaptic competition is induced by patterns of synaptic stimulation that induce robust homosynaptic LTP). Why might this be important? Because TPS-induces induces a Hebbian form of homosynaptic potentiation (Thomas et al., 1998; O’Dell, 2022), CS burst-dependent LTP likely shares a well-characterized, but problematic, feature of standard Hebbian plasticity rules – runaway dynamics. Runaway changes in synaptic strength in neural networks with Hebbian synapses are generated by the correlational nature of Hebbian plasticity rules, i.e., synapses potentiate in an unsupervised fashion in response to coincident pre- and postsynaptic activity. Thus, as some synapses undergo LTP, the increase in postsynaptic activity produced by these inputs increases the probability that other synapses undergo LTP. If left unopposed, this property of Hebbian LTP induction can trigger a self-reinforcing, positive-feedback cycle of potentiation that, as it continues, leads to runaway synaptic potentiation and loss of information encoded by differences in the strength of individual synapses (Miller and MacKay, 1994; Turrigiano, 2008; Zenke et al., 2013; Chen et al., 2013). Because of this, information storage in neural networks with Hebbian synapses requires rapid, negative-feedback forms of homeostatic plasticity to oppose the runaway dynamics of Hebbian LTP (Chistiakova et al., 2015; Zenke and Gerstner, 2017; Zenke et al., 2017). In CS burst-dependent LTP, this negative feedback is likely provided, at least in part, by the strong heterosynaptic suppression of LTP induction that emerges during bouts of EPSP-evoked CS bursting that induce robust homosynaptic LTP (O’Dell, 2022). Thus, the leftward shift in the inverted-U shaped relationship between S1 TPS duration and LTP induction at S2 synapses induced by ISO (Fig. 12B) suggests that β-AR activation not only enhances LTP induction but also recruits a compensatory increase in synaptic competition that can oppose the runaway dynamics of facilitated CS burst-dependent LTP induction.

CS bursts not only provide the postsynaptic depolarization needed for NMDAR activation and LTP induction during TPS (Thomas et al., 1998; O’Dell, 2022) but, as shown here, also trigger an A1 adenosine receptor-mediated heterosynaptic depression that underlies synaptic competition (Figs. 6 and 7). This dual role of CS bursts in LTP induction and synaptic competition thus provides a surprisingly simple, but computationally efficient, mechanism whereby increases in CS bursting triggered by β-AR activation can facilitate LTP induction while simultaneously ensuring sufficient synaptic competition is available to oppose runaway synaptic dynamics. The shared dependence of TPS-induced LTP and synaptic competition on postsynaptic CS bursts has other potentially important computational implications. For example, biologically realistic artificial neural networks with burst-dependent synaptic plasticity rules can implement the error back-propagation algorithm (Payeur et al., 2021; Sun et al., 2021), an especially powerful mathematical algorithm used in machine learning (Rumelhart et al., 1986). However, to successfully learn via error backpropagation these networks require a novel form of synaptic competition regulated by the probability of postsynaptic bursting, rather than the firing rate of single action potentials (Payeur et al., 2021). Although the properties of synaptic competition in CS burst-dependent LTP have not yet been fully characterized, the dual role of postsynaptic CS bursts in LTP induction and synaptic competition described here provide some initial experimental support for this theoretical prediction.

β-AR signaling at excitatory synapses in the CA1 region of the hippocampus is surprisingly complex (Jami et al., 2023). Thus, the identification of the molecular mechanisms underlying the β-AR modulation of synaptic interactions in CS burst-dependent LTP will require further investigation. The amount of homosynaptic LTP induced by different patterns of TPS is, however, highly correlated with the probability of EPSP-evoked CS bursting during TPS (O’Dell, 2022), as is the magnitude of TPS-induced heterosynaptic depression and synaptic competition (Fig. 4C,D). Similarly, there is a significant correlation between the heterosynaptic facilitation of EPSP-evoked CS bursting and cooperative LTP induction (Fig. 11E). Together, these findings suggest that β-AR-mediated increases in CA1 pyramidal cell excitability may have an especially important role in regulating CS burst-dependent LTP. Importantly, dendritic spikes and CS bursting in CA1 pyramidal cells are strongly attenuated by dendritic Kv4.2/A-type potassium channels (Magee and Carruth; 1999; Hoffman et al., 1997) and Kv1.1-type potassium channels inhibit action potential backpropagation in CA1 pyramidal cell dendrites (Liu et al., 2017). Thus, the inhibition of Kv4.2 (Hoffman and Johnston, 1999; Yuan et al., 2002) and Kv1.1 channels (Liu et al., 2017) induced by β-AR activation may have a key role in facilitating synaptic interactions during the induction of CS burst-dependent LTP. Because CS bursts in CA1 pyramidal cells are generated, in part, by activation of voltage-dependent calcium channels (Takahashi and Magee, 2009; Grienberger et al., 2014), β-AR-mediated increases in the activity of L-type calcium channels (Patriarchi et al., 2016; Qian et al., 2017) may also have an important role. Consistent with the notion that that changes in pyramidal cell excitability underlie the multiple effects of β-AR activation on CS burst-dependent LTP, the increase in EPSP-evoked CS bursting induced by the GABA_A_ receptor antagonist gabazine not only facilitates homosynaptic LTP induction but, like ISO, enhances TPS-induced heterosynaptic depression (Fig. 5) and increases synaptic competition (Fig. 6). However, β-AR activation had no effect on EPSP-evoked CS bursting during 5 seconds of TPS but nonetheless enabled the induction of LTP during these brief trains of TPS (Fig. 3A-C). This indicates that increases in CA1 pyramidal cell excitability cannot account for all the effects of β-AR activation on CS burst-dependent LTP. Thus, other mechanisms implicated in the β-AR modulation of LTP induction, such as changes in AMPA receptor trafficking (Hu et al., 2007; Qian et al., 2012), increases in NMDAR activity (Raman et al., 1996; Murphy et al., 2014), and/or inhibition of protein phosphatase signaling (Thomas et al., 1996), likely also contribute to the effects of β-AR activation on CS burst-dependent LTP (for review, see O’Dell et al., 2015).

Although the ability of β-AR activation to fundamentally alter the temporal and activity-dependent properties of synaptic interactions in CS burst-dependent plasticity is striking, there are several important questions to address in future experiments. First, bath application of modulatory neurotransmitter receptor agonists and release of endogenous transmitters can sometimes have very different effects on neuronal excitability and synaptic transmission (Rosen et al., 2015; Teixeira et al., 2018; Bacon et al., 2020). Although these findings raise concerns about the use of bath-applied receptor agonists to study the effects on modulatory neurotransmitters on synaptic plasticity, exogenous β-AR receptor agonists and optogenetic stimulation of locus coeruleus neurons have similar effects on dendritic excitability (Liu et al., 2017) and EPSP-spike coupling (Bacon et al., 2020) in CA1 pyramidal cells. ISO also mimics the effects of optogenetic activation of locus coeruleus axons on LTP induction at cortical synapses (He et al., 2015). Thus, exogenous β-AR agonists mimic the effects of endogenous NE on cellular processes likely to have an important role in CS bust-dependent LTP. Nevertheless, the use of optogenetic activation of locus coeruleus neurons will be important in future studies examining the noradrenergic regulation of CS burst-dependent LTP. Second, the facilitation of both memory formation and LTP induction induced by optogenetic activation of locus coeruleus fibers in the hippocampus may be mediated by the release of dopamine, rather than NE (Takeuchi et al., 2016; Wilmot et al., 2023, however, see Tsetsenis et al., 2022). Thus, a consideration of the role of dopamine receptor signaling will be important in future studies examining how modulatory inputs from the locus coeruleus regulate CS burst-dependent LTP. Finally, like BTSP, CS burst-dependent LTP exhibits a retroactive form of synaptic cooperativity where CS bursts can trigger LTP induction at synapses that were active seconds earlier (O’Dell, 2022). This form of cooperativity is thought to arise from the generation of synaptic eligibility traces, transient biochemical changes at synapses that, while having no direct effect on synaptic strength, persist for several seconds and enable synaptic potentiation in response to dendritic spikes induced by other synaptic inputs (Magee and Grienberger, 2020). Notably, β-AR activation can induce LTP in a retrograde fashion at cortical synapses by transforming eligibility traces generated by prior synaptic activity (He et al., 2015). Importantly, by providing a mechanism that allows temporally delayed signals related to prediction errors or reward to induce changes in synaptic strength, eligibility traces potentially solve the long-standing “distal reward problem” in reinforcement learning (Izhikevich, 2007; Magee and Grienberger, 2020). Thus, it will be interesting to determine whether a β-AR-mediated transformation of TPS-induced eligibility traces also contribute to the surprisingly complex effects of β-AR activation on the properties of CS burst-dependent LTP.

## Acknowledgements

This work was supported by funds provided by the UCLA Academic Senate. Funders had no role in study design, data collection/analysis, manuscript preparation, or publication.

---

**Supplemental Figure 1.**
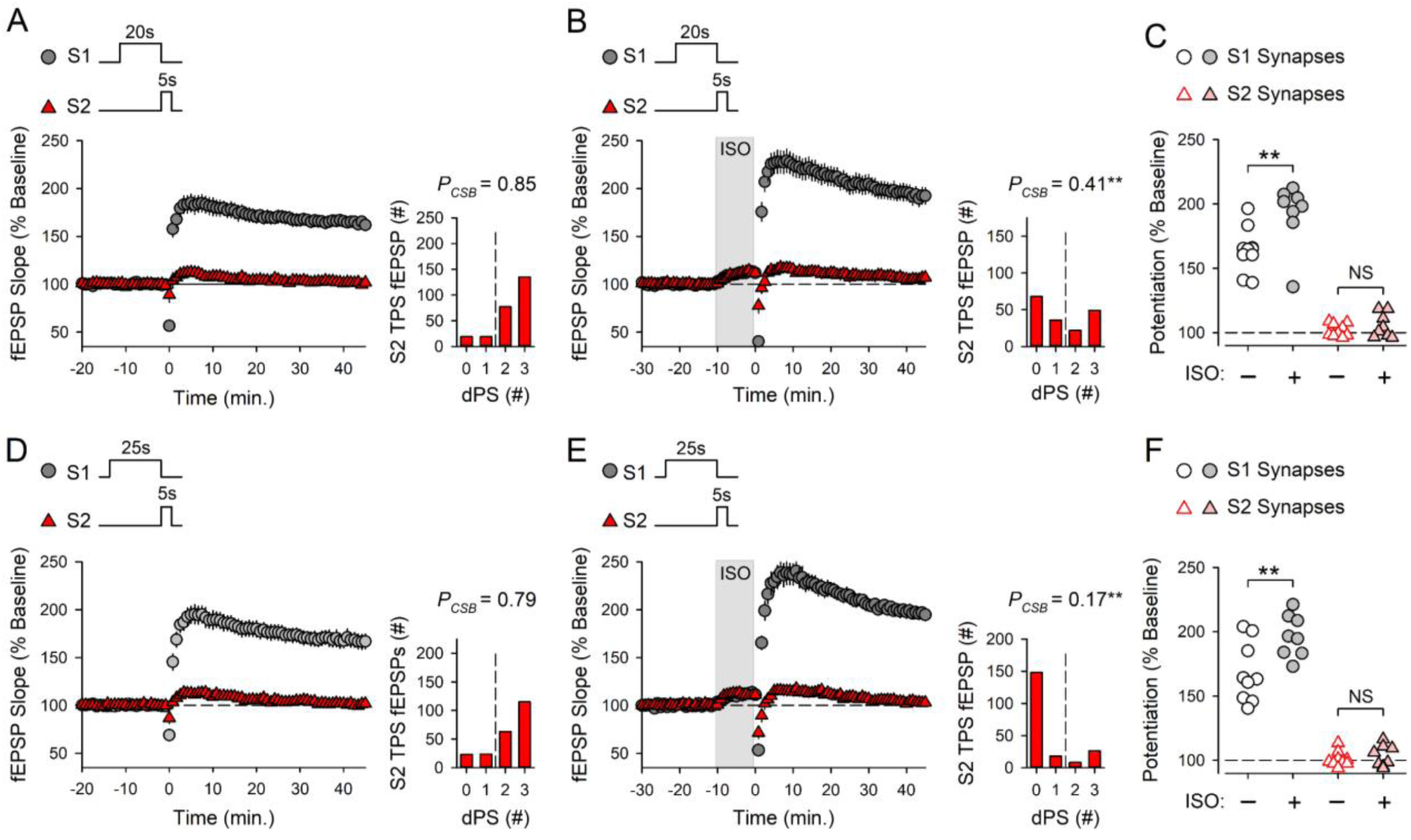
β-AR activation enhances the induction of LTP at S1 synapses but has no effect on the suppression of LTP induction at S2 synapses induced by long trains of S1 TPS. ***A*:** Control experiments where 20 seconds of TPS delivered to S1 synapses before 5 seconds of S2 TPS (at time = 0). 45 minutes post-TPS S1 synapses potentiated to 164 ± 5% of baseline and S2 synapses were 101 ± 2% of baseline (n = 10). ***B*:** 20 seconds of S1 TPS before S2 TPS was delivered at the end of a 10-minute bath application of 1.0 µM ISO (indicated by the shaded region). 45 minutes post-TPS S1 were 192 ± 9% of baseline and S2 synapses were 105 ± 3% of baseline (n = 8). **C:** S1 and S2 fEPSP slopes 45 minutes post-S2 TPS from all experiments in *A* and *B*. Results were analyzed a using two-way ANOVA with SNK *post hoc* comparisons (\*\**p* < 0.001, NS, not significant, *p* = 0.538). There was a significant difference between S1 and S2 synapses (*F*_(1,32)_ = 208.672, *p* < 0.001), a significant effect of ISO (*F*_(1,32)_ = 9.779, *p* = 0.004), and a significant synapse x ISO interaction (*F*_(1,32)_ = 5.490, *p* = 0.025). ***D*:** Control experiments where 25 seconds of S1 TPS was delivered before 5 seconds of S2 TPS. 45 minutes post-TPS S1 synapses were 168 ± 8% of baseline and S2 synapses were 101 ± 2% of baseline (n = 9). ***E*:** 25 seconds of S1 TPS before S2 TPS was delivered in the presence of ISO. 45 minutes post-TPS S1 synapses were 197 ± 6% of baseline and S2 synapses were 103 ± 3% of baseline (n = 8). ***F*:** S1 and S2 fEPSP slopes 45 minutes post-S2 TPS from all experiments in *D* and *E*. A two-way ANOVA with SNK post hoc comparisons (\*\**p* < 0.001, NS, not significant, *p* = 0.738) revealed a significant difference between S1 and S2 synapses (*F*(1,30) = 232.361, *p* < 0.001), a significant effect of ISO (*F*_(1,30)_ = 8.750, *p* = 0.006), and a significant synapse x ISO interaction (*F*_(1,30)_ = 6.152, *p* = 0.019). Histograms show number of EPSPs evoking 0 – 3 CS bursts during S2 TPS in all experiments. EPSP-evoked CS bursting during S2 TPS was significantly reduced by ISO (20s S1 TPS: Mann Whitney *U* = 12.5, \**p* = 0.019, 25s S1 TPS: Mann Whitney U = 11.5, \**p* = 0.017). This is likely due to the strong heterosynaptic depression induced by 20 and 25 seconds of S1 TPS in the presence of ISO (fEPSPs evoked during S2 TPS were reduced to ∼20% of baseline, data not shown).

## REFERENCES

Babiec, W.E., S.A. Jami, R., R. Guglietta, P.B. Chen, and T.J. O’Dell. 2017. Differential Regulation of NMDA Receptor-Mediated Transmission by SK Channels Underlies Dorsal-Ventral Differences in Dynamics of Schaffer Collateral Synaptic Function. Journal of Neuroscience 37: 1950–1964. 10.1523/JNEUROSCI.3196-16.2017

Bacon, T.J., A.E. Pickering, and J.R. Mellor JR. 2020. Noradrenaline Release from Locus Coeruleus Terminals in the Hippocampus Enhances Excitation-Spike Coupling in CA1 Pyramidal Neurons via Beta-Adrenoceptors. Cerebral Cortex 30: 6135–6151. 10.1093/cercor/bhaa159

Bittner, K.C., A.D. Milstein, C. Grienberger, S. Romani, and J.C. Magee JC. 2017. Behavioral Time Scale Synaptic Plasticity Underlies CA1 Place Fields. Science 357: 1033–1036. 10.1126/science.aan3846.

Bouret, S. and S.J. Sara. 2004. Reward Expectation, Orientation of Attention and Locus Coeruleus-Medial Frontal Cortex Interplay During Learning. European Journal of Neuroscience 20: 791–802. 10.1111/j.1460-9568.2004.03526.x.

Brzosko, Z., S.B. Mierau, and O. Paulsen. 2019. Neuromodulation of Spike-Timing-Dependent Plasticity: Past, Present, and Future. Neuron 103: 563–581. 10.1016/j.neuron.2019.05.041.

Chen, J.-Y., P. Lonjers, C. Lee, M. Chistiakova, M. Volgushev, and M. Bazhenov. 2013. Heterosynaptic Plasticity Prevents Runaway Synaptic Dynamics. Journal of Neuroscience 33: 15915–15929. 10.1523/JNEUROSCI.5088-12.2013

Chistiakova, M., N.M. Bannon, J.-Y. Chen, M. Bazhenov, and M. Volgushev. 2015. Homeostatic Role of Heterosynaptic Plasticity: Models and Experiments. Frontiers in Computational Neuroscience 9: 89. 10.3389/fncom.2015.00089

Dringenberg, H.C. 2019. The History of Long-Term Potentiation as a Memory Mechanism: Controversies, Confirmation, and Some Lessons to Remember. Hippocampus 30: 987–1012. 10.1002/hipo.23213.

Fanselow, M.S., and K.M. Wassum. 2016. The Origins and Organization of Vertebrate Pavlovian Conditioning. Cold Spring Harbor Perspectives in Biology 8: a021717. https://cshperspectives.cshlp.org/content/8/1/a021717.

Frémaux, N., and W. Gerstner. 2016. Neuromodulated Spike-Timing-Dependent Plasticity, and Theory of Three-Factor Learning Rules. Frontiers in Neural Circuits 9: 85. 10.3389/fncir.2015.00085.

Gallistel, C.R., and L.D. Matzel. 2013.The Neuroscience of Learning: Beyond the Hebbian Synapse. Annual Review of Psychology 64: 169–200. 10.1146/annurev-psych-113011-143807.

Gelinas, J.N., and P.V. Nguyen. 2005. β-Adrenergic Receptor Activation Facilitates Induction of a Protein Synthesis-Dependent Late Phase of Long-Term Potentiation. Journal of Neuroscience 25: 3294–3303. 10.1523/JNEUROSCI.4175-04.2005.

Gelinas, J.N., G. Tenorio, N. Lemon, T. Abel, and P.V. Nguyen. 2008. β-Adrenergic Receptor Activation During Distinct Patterns of Stimulation Critically Modulates the PKA-Dependence of LTP in the Mouse Hippocampus. Learning and Memory 15: 281–289. http://www.learnmem.org/cgi/doi/10.1101/lm.829208.

Grienberger, C., X. Chen, and A. Konnerth. 2014. NMDA Receptor-Dependent Multidendrite Ca^2+^ Spikes Required for Hippocampal Burst Firing In-Vivo. Neuron 81: 1274–1281. 10.1016/j.neuron.2014.01.014.

Grienberger, C., and J.C. Magee. 2022. Entorhinal Cortex Directs Learning-Related Changes in CA1 Representations. Nature 611: 554–562. 10.1038/s41586-022-05378-6.

Grover, L.M., and T.J. Teyler. 1993. Role of Adenosine in Heterosynaptic, Posttetanic Depression in Area CA1 of Hippocampus. Neuroscience Letters 154: 39–42. 10.1016/0304-3940(93)90166-I

Gustafsson, B., and H. Wigström. 1986. Hippocampal Long-Lasting Potentiation Produced by Pairing Single Volleys and Brief Conditioning Tetani Evoked in Separate Afferents. Journal of Neuroscience 6: 1575–1582. 10.1523/JNEUROSCI.06-06-01575.1986.

Hagena, H., N. Hansen, and D. Manahan-Vaughan. 2016. β-Adrenergic Control of Hippocampal Function: Subserving the Choreography of Synaptic Information Storage and Memory. Cerebral Cortex 26: 1349–1364. 10.1093/cercor/bhv330.

Hoffman, D.A., and D. Johnston D. 1999. Neuromodulation of Dendritic Action Potentials. Journal of Neurophysiology 81: 408–411. 10.1152/jn.1999.81.1.408

Hoffman, D.A., J.C. Magee, C.M. Colbert CM, and D. Johnston. 1997. K^+^ Channel Regulation of Signal Propagation in Dendrites of Hippocampal Pyramidal Neurons. Nature 387: 869–875. 10.1038/43119

He, K., M. Huertas, S.Z. Hong, X.X. Tie, J.W. Hell, H. Shouval, and A. Kirkwood. 2015. Distinct Eligibility Traces for LTP and LTD in Cortical Synapses. Neuron 88: 528–538. 10.1016/j.neuron.2015.09.037

Hu, H., E. Real, K. Takamiya, M.-G. Kang, J. Ledoux, R.L. Huganir, and R. Malinow. 2007. Emotion Enhances Learning via Norepinephrine Regulation of AMPA-Receptor Trafficking. Cell 131: 160–173. 10.1016/j.cell.2007.09.017

Izhikevich, E.M. 2007. Solving the Distal Reward Problem Through Linkage of STDP and Dopamine Signaling. Cerebral Cortex 17: 2443–2452. 10.1093/cercor/bhl152

Jami, S.A., B.J. Wilkinson, R. Guglietta, N. Hartel, W.E. Babiec, N.A. Graham, M.P. Coba, and T.J. O’Dell. 2023. Functional and phosphoproteomic analysis of β-adrenergic receptor signaling at excitatory synapses in the CA1 region of the ventral hippocampus. Scientific Reports 13: 7493. 10.1038/s41598-023-34401-7.

Jorden, R. 2024. The Locus Coeruleus as a Global Model Failure System. Trends in Neuroscience 47: 92–105. 10.1016/j.tins.2023.11.006.

Kaufman, A.M., T. Geiller, and A. Losonczy. 2022. A Role for the Locus Coeruleus in Hippocampal CA1 Place Cell Reorganization during Spatial Reward Learning. Neuron 105: 1018–1026. 10.1016/j.neuron.2019.12.029.

Lin, Y.-W., M.-Y. Min, T.-H. Chiu, and H.-W. Yang. 2003. Enhancement of Associative Long-Term Potentiation by Activation of β-Adrenergic Receptors at CA1 Synapses in Rat Hippocampal Slices. Journal of Neuroscience 23: 4173–4181. 10.1523/JNEUROSCI.23-10-04173.2003.

Liu, Y., L. Cui, M.K. Schwarz, Y. Don, and O.M. Schlüter. 2017. Adrenergic Gate Release for Spike Timing-Dependent Synaptic Potentiation. Neuron 93: 394–408. 10.1016/j.neuron.2016.12.039

Lovatt, D., Q. Xu, W. Liu, T. Takano, N.A. Smith, J. Schnermann, K. Tieu, and M. Nedergaard. 2012. Neuronal Adenosine Release, and not Astrocytic ATP Release, Mediates Feedback Inhibition of Excitatory Activity. Proceedings of the National Academy of Sciences 109: 6265–6270. 10.1073/pnas.1120997109

Lovett-Barron, M., G.F. Turi, P. Kaifosh, P.H. Lee, F. Bolze, X.-H. Sun, J.-F. Nicoud, B.V. Zemelman, S.M. Sternson, and A. Losonczy. 2012. Regulation of Neuronal Input Transformation by Tunable Dendritic Inhibition. Nature Neuroscience 15: 423–430. 10.1038/nn.3024.

Magee, J.C., and M. Carruth 1999. Dendritic Voltage-Gated Ion Channels Regulate the Action Potential Firing Mode of Hippocampal CA1 Pyramidal Cells. Journal of Neurophysiology 83: 1895–1901. 10.1152/jn.1999.82.4.1895

Magee, J.C., and C. Grienberger. 2020. Synaptic Plasticity Forms and Functions. Annual Review of Neuroscience 43: 95–117. 10.1146/annurev-neuro-090919-022842

Miller, K.D., and D.J.C MacKay. 1994. The Role of Constraints in Hebbian Learning. Neural Computation 6: 100–126. 10.1162/neco.1994.6.1.100

Mitchell, J.B., C.R. Lupica, and T.V. Dunwiddie. 1993. Activity-Dependent Release of Endogenous Adenosine Modulates Synaptic Responses in the Rat Hippocampus. Journal of Neuroscience 13: 3439–3447. 10.1523/JNEUROSCI.13-08-03439

Murphy, J.A., I.S. Stein, C.G. Lau, R.T. Peixoto, T.K. Aman, N. Kaneko, K. Aromolaran, J. L. Saulnier, G.K. Popescu, B.L. Sabatini, J.W. Hell, and R.S. Zukin. 2014. Phosphorylation of Ser1166 on GluN2B by PKA is Critical to Synaptic NDMA Receptor Function and Ca^2+^ Signaling in Spines. Journal of Neuroscience 34: 869–879. 10.1523/JNEUROSCI.4538-13.2014

O’Dell, TJ. 2022. Behavioral Timescale Cooperativity and Competitive Synaptic Interactions Regulate the Induction of Complex Spike Burst-Dependent Long-Term Potentiation. Journal of Neuroscience 42: 2647–2661. 10.1523/JNEUROSCI.1950-21.2022.

O’Dell, T.J., S.A. Connor, R. Guglietta, and P.V. Nguyen. 2015. β-Adrenergic Receptor Signaling and Modulation of Long-Term Potentiation in the Mammalian Hippocampus. Learning and Memory 22: 461–471. http://www.learnmem.org/cgi/doi/10.1101/lm.031088.113.

Patriarchi, T., H. Qian, V. Di Biase, Z.A. Malik, D. Chowdhury, J.L. Price, E.A. Hammes, O.R. Buonarati, R.E. Westenbroek, W.A. Catterall, F. Hofmann, Y.K. Xiang, G.G. Murphy, C.-Y. Chen, M.F. Navedo, and J.W. Hell. 2016. Phosphorylation of Cav1. 2 on S1928 Uncouples the L-Type Ca^2+^ Channel from the β2 Adrenergic Receptor. The EMBO Journal 35, 1330–1345. 10.15252/embj.201593409

Payeur, A., J. Guerguiev, F. Zenke, B.A. Richards, and R. Naud. 2021. Burst-Dependent Synaptic Plasticity can Coordinate Learning in Hierarchical Circuits. Nature Neuroscience 24: 1010–1019. 10.1038/s41593-021-00970-x

Priestley, J.B., J.C. Bowler, S.V. Rolotti, S. Fusi, and A. Losonczy. 2022. Signatures of Rapid Plasticity in Hippocampal CA1 Representations During Novel Experiences. Neuron 110: 1978–1992. 10.1016/j.neuron.2022.03.026.

Qian, H., L. Matt, M. Zhang, M. Nguyen, T. Patriarchi, O.M. Koval, M.E. Anderson, K. He, H.-K. Lee, and J.W. Hell. 2012. β2-Adrenergic Receptor Supports Prolonged Theta Tetanus-Induced LTP. Journal of Neurophysiology 107: 2703–2712. 10.1152/jn.00374.2011.

Qian, H., T. Patriarchi, J.L. Prince, L. Matt, B. Lee, M. Nieves-Cintrón, O.R. Buonarati, D. Chowdhury, E. Nanou, M.A. Nystoriak, W.A. Catterall, M. Poomvanicha, F. Hofmann, M.F. Navedo, and J.W. Hell. 2017. Phosphorylation of Ser1928 Mediates the Enhanced Activity of the L-Type Ca^2+^ Channel Cav1.2 by the β2-Adrenergic Receptor in Neurons. Science Signaling 10, eaaf9659. 10.1126/scisignal.aaf9659

Raman, I.M., G. Tong, and C.E. Jahr. 1996. β-adrenergic regulation of synaptic NMDA receptors by cAMP-dependent protein kinase. Neuron 16: 415–421. 10.1016/S0896-6273(00)80059-8

Rumelhart, D.E., G.E. Hinton, and R.J. Williams. 1986. Learning Representations by Back-Propagating Errors. Nature 323: 533–536. 10.1038/323533a0

Rosen, Z.B., S. Cheung, and S.A. Siegelbaum. 2015. Midbrain Dopamine Neuros Bidirectionally Regulate CA3-CA1 Synaptic Drive. Nature Neuroscience 18: 1763–1771. 10.1038/nn.4152

Sara, S.J. 2009. The Locus Coeruleus and Noradrenergic Modulation of Cognition. Nature Reviews Neuroscience 10: 211–223. 10.1038/nrn2573.

Sara, S.J., A. Vankov, and A. Hervé. 1994. Locus Coeruleus-Evoked Responses in Behaving Rats: A Clue to the Role of Noradrenaline in Memory. Brain Research Bulletin 35: 457–464. 10.1016/0361-9230(94)90159-7.

Sun, W., X. Zhao, and N. Spruston. 2021. Bursting Potentiates the Neuro-AI Connection. Nature Neuroscience 24: 905–906. 10.1038/s41593-021-00844-2

Takahashi, H., and J.C. Magee. 2009. Pathway Interactions and Synaptic Plasticity in the Dendritic Tuft Regions of CA1 Pyramidal Neurons. Neuron 62: 102–111. 10.1016/j.neuron.2009.03.007.

Takeuchi, T., A.J. Duszkiewicz, A. Sonneborn, P.A. Spooner, M. Yamasaki, M. Watanabe, C.C. Smith, G. Fernandez, K. Deisseroth, R.W. Greene, and R.G. Morris. 2016. Locus Coeruleus and Dopaminergic Consolidation of Everyday Memory. Nature 37: 357–362. 10.1038/nature19325.

Teixeira, C.M., Z.B. Rosen, D. Suri, Q. Sun, M. Hersh, D. Sargin, I. Dincheva, A. Morgan, S. Spivack, A.C. Krok, T. Hirschfeld-Stoler, E.K. Lambe, S.A. Siegelbaum, and M.S. Ansorge 2018. Hippocampal 5-HT Input Regulates Memory Formation and Schaffer Collateral Excitation. Neuron 98: 992–1004. 10.1016/j.neuron.2018.04.030

Thomas, M.J., T.D. Moody, M. Makhinson, and T.J. O’Dell. 1996. Activity-Dependent β-Adrenergic Modulation of Low-Frequency Stimulation Induced LTP in the Hippocampal CA1 Region. Neuron 17: 475–482. 10.1016/S0896-6273(00)80179-8.

Thomas, M.J., A.M. Watabe, T.D. Moody, M. Makhinson, and T.J. O’Dell. 1998. Postsynaptic Complex Spike Bursting Enables the Induction of LTP by Theta Frequency Synaptic Stimulation. Journal of Neuroscience 18: 7118–7126. 10.1523/JNEUROSCI.18-18-07118.1998.

Tsetsenis, T., J.K. Badyna, R. Li, and J.A. Dani. 2022. Activation of a Locus Coeruleus to Dorsal Hippocampus Noradrenergic Circuit Facilitates Associative Learning. Frontiers in Cellular Neuroscience 16: 887679. 10.3389/fncel.2022.887679

Turrigiano. G.G. 2008. The Self-Tuning Neuron: Synaptic Scaling of Excitatory Synapses. Cell 135: 422–435. 10.1016/j.cell.2008.10.008

Wigström, H., and B. Gustafsson. 1984. Facilitated Induction of Hippocampal Long-Lasting Potentiation During Blockade of Inhibition. Nature 301: 603–604. 10.1038/301603a0

Wilmot, J.H., C.R.A.F. Diniz, A.P. Crestani, K.R. Puhger, J. Roshgadol, L. Tian, and B.J. Wiltgen. 2023. Phasic Locus Coeruleus Activity Enhances Trace Fear Conditioning by Increasing Dopamine Release in the Hippocampus. eLife 12: RP91465. 10.7554/eLife.91465

Winder, D.G., K.C. Martin, I.A. Muzio, D. Rohrer, A. Chruscinski, B. Kobilka, and E.R. Kandel. 1999. ERK Plays a Regulatory Role in Induction of LTP by Theta Frequency Stimulation and its Modulation by β-Adrenergic Receptors. Neuron 24: 715–726. 10.1016/S0896-6273(00)81124-1.

Wu, L.-G., and P. Saggau. 1994. Adenosine Inhibits Evoked Synaptic Transmission Primarily by Reducing Presynaptic Calcium Influx in Area CA1 of Hippocampus. Neuron 12: 1139–1148. 10.1016/0896-6273(94)90321-2

Wu, Z., Y. Cui, H. Wang, H. Wu, Y. Wan, B. Li, L. Wang, S. Pan, W. Peng, A. Dong, Z. Yuan, M. Jing, M. Zu, M. Luo, and Y. Li. 2023. Neuronal Activity-Induced, Equilibrative Nucleoside Transporter-Dependent, Somatodendritic Adenosine Release Revealed by a GRAB Sensor. Proceedings of the National Academy of Sciences 120: e2212387120. 10.1073/pnas.2212387120

Xiao, K., Y. Li, R.A. Chitwood, and J.C. Magee. 2023. A Critical Role for CaMKII in Behavioral Timescale Synaptic Plasticity in Hippocampal CA1 Pyramidal Cells. Science Advances 9: eadi3088. 10.1126/sciadv.adi3088.

Yuan, L.-L., J.P. Adams, M. Swank, J.D. Sweatt, and D. Johnston. 2002. Protein Kinase Modulation of Dendritic K^+^ Channels in Hippocampus involves a Mitogen-Activated Protein Kinase Pathway. Journal of Neuroscience 22: 4860–4868. 10.1523/JNEUROSCI.22-12-04860.2002

Zhao, X., C.-L. Hsu, and N. Spruston. 2022. Rapid Synaptic Plasticity Contributes to a Learned Conjunctive Code of Position and Choice-Related Information in the Hippocampus. Neuron 110: 96–108. 10.1016/j.neuron.2021.10.003.

Zenke, F., and W. Gerstner. 2017. Hebbian Plasticity Requires Compensatory Processes on Multiple Timescales. Philosophical Transactions of the Royal Society B 372: 20160259. 10.1098/rstb.2016.0259

Zenke, F., W. Gerstner, and S. Ganguli. 2017. The Temporal Paradox of Hebbian Learning and Homeostatic Plasticity. Current Opinion in Neurobiology 43: 166–176. 10.1016/j.conb.2017.03.015

Zenke, F., G. Hennequin, and W. Gerstner. 2013. Synaptic Plasticity in Neural Networks Needs Homeostasis with a Fast Rate Detector. PLoS Computational Biology 9(11): e1003330. 10.1371/journal.pcbi.1003330

